# SWI/SNF Alterations Define a Chromatin-Dependent Subtype of Urothelial Carcinoma

**DOI:** 10.64898/2026.08.05.743022

**Authors:** Bing-Jian Feng, Kaniz Fatema, David A. Nix, Aaron Atkinson, Constance Caparas, Chris J. Stubben, David Henry Lum, Timothy J. Parnell, Courtney Carroll, G. Daniel Grass, Laura Graham, Eric A. Singer, Kenneth G. Nepple, Zarko Manojlovic, Eric Kauffman, Jennifer M. King, Saum Ghodoussipour, Patrick Hensley, Paul V. Viscuse, Adanma Ayanambakkam, Michelle L. Churchman, Umang Swami, Neeraj Agarwal, Bradley R. Cairns, Sumati Gupta

**Affiliations:** Department of Dermatology, University of Utah, 30 N. Mario Capecchi Dr., Salt Lake City, UT, 84112, USA; Huntsman Cancer Institute, University of Utah, 2000 Circle of Hope Drive, Salt Lake City, UT, 84112, USA; Moffitt Cancer Center; University of Colorado Anschutz; Division of Urologic Oncology, The Ohio State University Comprehensive Cancer Center, Columbus, OH, USA; University of Iowa Holden Comprehensive Cancer Center; Department of Urology, University of Southern California, Los Angeles, CA, USA; Roswell Park Comprehensive Cancer Center; Indiana University School of Medicine; Cancer Institute of New Jersey; Markey Cancer Center; University of Virginia Comprehensive Cancer Center; University of Oklahoma Health; Aster Insights

## Abstract

**Purpose:** SWI/SNF (BAF) chromatin remodeling complex alterations are common in urothelial carcinoma, yet no biomarker-directed therapeutic strategies have been established for this population. We investigated whether BAF alterations delineate a biologically distinct, therapeutically actionable urothelial carcinoma subtype.

**Experimental Design:** We performed integrative genomic and transcriptomic analyses of 792 urothelial carcinoma tumors from the Oncology Research Information Exchange Network (ORIEN) and validated findings in the TCGA-BLCA cohort. Mechanistic studies incorporated RNA sequencing and ATAC-seq following histone deacetylase (HDAC) inhibition. Functional dependencies were assessed using patient-derived xenograft organoids and cell line models. Clinical relevance was explored in a biomarker-enriched investigator-initiated trial.

**Results:** Approximately half of urothelial carcinoma tumors exhibited BAF alterations, defining a previously unrecognized chromatin-altered molecular subtype characterized by activation of proliferative programs, loss of lineage identity, and altered metabolic signaling. This subtype was enriched for transcriptomic programs associated with HDAC inhibitor sensitivity and depleted of HDAC inhibitor resistance signatures. Mechanistically, HDAC inhibition induced widespread chromatin remodeling with reduced accessibility at AP-1 and TEAD-associated regions, and downregulation of E2F- and MYC-driven transcriptional networks. Functional studies confirmed enhanced HDAC inhibition sensitivity in *ARID1A*-mutated cell lines and a patient-derived organoid model. Early clinical observations demonstrated a durable responder treated with HDAC inhibitors and immunotherapy.

**Conclusions:** BAF alterations define a chromatin-dependent tumor state in urothelial carcinoma that is selectively vulnerable to HDAC inhibition. Integrating genomic, epigenomic, functional, and early clinical evidence, these findings provide a rationale for biomarker-enriched clinical trials and HDAC inhibitor-based combination strategies in urothelial carcinoma.

## INTRODUCTION

Urothelial carcinoma is a clinically and molecularly heterogeneous disease for which biomarker-directed therapeutic strategies remain limited. While recent advances in immune checkpoint inhibition and antibody–drug conjugates have improved outcomes in patients with urothelial carcinoma, many treatment decisions are not guided by tumor genomics. Identifying molecularly defined populations with actionable therapeutic vulnerabilities remains a critical unmet need.

The switch/sucrose nonfermenting-type (SWI/SNF) chromatin remodeling complexes, also referred to as BRG1/BRM-associated factor (BAF) complexes, are multiprotein complexes that regulate the accessibility of genomic DNA to transcription factors.^1^ SWI/SNF alterations typically lead to loss of tumor suppressor gene function, which broadly alters chromatin accessibility and transcription, making it challenging to directly target these alterations.^2^

SWI/SNF alterations are highly prevalent in urothelial carcinoma, occurring in more than half of tumors, yet have not been successfully translated into predictive biomarkers or therapeutic strategies. It remains unknown whether mutations in distinct BAF subunits converge on a unified therapeutic vulnerability. We hypothesized that SWI/SNF alterations define a biologically distinct and therapeutically targetable subtype of urothelial carcinoma characterized by a convergent transcriptional and chromatin-defined state. We leveraged a dataset of whole-exome and RNA sequencing data from 792 patients with urothelial carcinoma in the Oncology Research Information Exchange Network (ORIEN) to test our hypothesis and validated key findings in the Cancer Genome Atlas - Bladder urothelial carcinoma cohort (TCGA-BLCA). We identified the transcriptional signature and therapeutic vulnerability of urothelial carcinoma with BAF alterations, with a translational focus on *ARID1A-*mutant urothelial carcinoma. We present preclinical functional validation and early clinical translation from the RESOLVE trial (NCT05154994).

## METHODS

### Ethics approval

This study was conducted in accordance with the Declaration of Helsinki and approved by the institutional review board.

### Analyses of ORIEN urothelial carcinoma data

The ORIEN Avatar program comprises 19 cancer centers that sequence the DNA and RNA from cancer tissue and the germline DNA of consenting patients to facilitate precision-based cancer clinical trials.^3^ ORIEN Avatar specimens were sequenced using previously described methods.^4^ The process and data sources are described on the ORIEN website at https://www.oriencancer.org. Bioinformatic analyses of the sequence and variant data have been previously described.^5,6^

Cancer driver genes were obtained from the Integrative OncoGenomics (IntOGen) database.^7^ Co-occurrence or mutual exclusivity between alterations in BAF genes and alterations in non-BAF genes were tested by the DISCOVER program.^8^

The TNRunner RNAAlignQC workflow was run on each tumor RNA-seq dataset. This workflow used CutAdapt 3.4 to remove adapter sequences, STAR 2.7.9a to align reads to the human GRCh38 Ensembl 106 reference, and featureCounts subread 2.0.3 to assign reads to annotated genes. To account for technical noise and latent biological variation in the ORIEN transcriptomic data, we applied Remove Unwanted Variation (RUVg) using the RUVSeq package. We identified 150 empirically stable genes across the dataset to serve as negative controls for the RUVg algorithm, setting the number of latent factors of unwanted variation (k) to 6. Following RUVg correction, the normalized counts were transformed with a variance stabilizing transformation in DESeq2 to produce a homoscedastic expression matrix for downstream visualization and pathway scoring.

Differential expression analysis was performed on 478 RNA-seq of primary tumors using limma with linear models adjusting for age, sex, stage, and estimated factors of unwanted variation.^9,10^ To further ensure the robustness of our results, the model was additionally adjusted for tissue type (Formalin-fixed paraffin-embedded (FFPE) vs. fresh frozen) and library preparation method (TruSeq vs. Tagmentation). Gene set enrichment analysis with Hallmark gene sets from the Molecular Signatures Database was subsequently performed with the CAMERA algorithm in the limma package.^11^

### Analysis of TCGA-BLCA data

The TCGA-BLCA WES tumor data were obtained from the cBioPortal. Variants were filtered and analyzed using the VICTOR package. Homozygous deletions were treated as truncating variants. RSEM (RNA-Seq by Expectation-Maximization) expected counts were used for differential gene expression analyses. Duplicate gene rows were collapsed by retaining the row with the lowest percentage of zero expression values; remaining ties were resolved by calculating the column-wise mean. Counts were then transformed to the log_2_(RSEM+1) scale. Low-expression genes were filtered out using a stringent multi-step thresholding approach, requiring genes to have an expression signal (log_2_>2) in at least 50% of the cohort and to maintain an overall mean expression greater than 1. Continuous cross-sample normalization was performed using quantile normalization via the limma package, and data heteroscedasticity and sample distributions were monitored using mean-variance trend plots (plotSA) and principal component analysis.

### 2D cell culture and Viability Assay

The HT-1197 (ATCC CRL-1473, RRID:CVCL_1291) and HT-1376 (ATCC CRL-1472, RRID:CVCL_1292) cell lines were obtained from ATCC and were cultured in ATCC-formulated Eagle’s Minimum Essential Medium (ATCC, Cat# 30-2003) and supplemented with 10% fetal bovine serum (FBS), 1% penicillin-streptomycin, and maintained at 37°C in a humidified incubator with 5% CO₂. Media were replaced every 2-4 days, and cells were passaged upon reaching 70–80% confluence.HT-1197 and HT-1376 cells (ATCC) were seeded in triplicate into 96-well plates at a density of 4,000 cells per well. 3-fold serial dilutions of belinostat (SelleckChem, S1085) were added starting at a concentration of 10 µM. After 72 hours of incubation at 37°C, cell viability was assessed using CellTiter-Glo (Promega). Half-maximal inhibitory concentration (IC₅₀) values were calculated and graphed using GraphPad Prism (version 9.4.1).

### ATAC-seq

ATAC-seq sample preparation was done using the Active Motif ATAC-Seq Kit following the included ‘Cell Sample Preparation’, ‘Tagmentation Reaction and Purification’, and ‘PCR Amplification of Tagmented DNA’ protocols. Three biological samples for each treatment (belinostat and control) were then submitted to the Huntsman Cancer Institute High-Throughput Genomics Core for paired-end sequencing on an Illumina Novaseq 6000. Reads were aligned to Hg38 using Novocraft Novoalign, v4.03.01. Peaks were called using the Multi-Replica Macs ChIPSeq Wrapper (https://huntsmancancerinstitute.github.io/MultiRepMacsChIPSeq) pipeline, version 17.9. Briefly, alignments were de-duplicated at a consistent 5% rate by subsampling duplicate alignments. Mean coverage tracks were generated by treating alignments as single-end, shifting the 5’ end by –25 bp and extending 50 bp, depth-normalizing each replicate individually, and then averaging across replicates. Peaks were called with a minimum q-value of 0.001. Called peaks from each condition were merged for joint analysis. Differential analysis was performed using DESeq2 with pre-normalized counts generated by the peak-calling pipeline. Motif analysis was performed using Homer v4.11 (http://homer.ucsd.edu/homer/) with the findMotifsGenome and annotatePeaks tools.

### RNA-seq

RNA-seq sample preparation was done using the Qiagen RNeasy kit and following the manufacturer’s protocol, including the ‘Purification of Total RNA from Animal Cells Using Spin Technology’ protocol. Three biological replicates from each condition were submitted to the Huntsman Cancer Institute High-Throughput Genomics Core for sequencing on an Illumina Novaseq6000. Reads were deduplicated using BBmap Clumpify (v38), adapters removed with Cutadapt (v3.5), aligned to GRCh38 using STAR (v2.7.6) and Ensembl annotation (release 104), and generated gene counts with Subread featureCounts. Differential genes were identified using DESeq2 with a default false discovery rate (FDR) cutoff of 0.01.

### Patient-derived Xenograft Organoid (PDxO) Culture

Organoids were generated from banked patient-derived xenograft (PDX) tumor chunks as previously described.^12,13^

PDxOs were cultured in the organoid growth medium optimized for specific bladder cancer organoids. The organoid culture medium was comprised of 50% basis media supplemented with 50% L-WRN (WNT3A, R-Spondin1, and Noggin-conditioned medium from HCI PRR core), 1.25 mM N-Acetyl-L-cysteine (Sigma-Aldrich, A9165), 10 μM Y-27632 ROCK inhibitor (Selleckchem, S6390), 50 ng/mL Epidermal Growth Factor (PeproTech, AF-100-15-1MG), 5% FBS, 1× N2 (GIBCO CTS, A1370701), 1× B27 solution (GIBCO, 17504044), and 1μM LY2109761 TGF-β receptor type I/II dual inhibitor (Selleck, S2704). Organoids were maintained in Ultra-Low Attachment (Corning® Costar®, CLS7007) culture dishes at 37°C in a humidified incubator with 5% CO₂. The medium was refreshed every 3-4 days, and cultures were passaged every two weeks. To confirm the genetic fidelity of organoids to their parental PDX tumors (internally designated as BLA-001), Short Tandem Repeat (STR) analysis was performed by HCI Preclinical Research Resources.

### PDxO Viability Assay

Organoids were collected and washed in the basis medium (100 g, 5 min) before being dissociated into single cells using 1 mL TrypLE Express and Y-27632 ROCK inhibitor (at a 1000:1 ratio) at 37°C for 10 minutes. Cells were vigorously pipetted every 2–3 minutes until a single-cell suspension was confirmed under the microscope. The suspension was then counted, washed once in basis medium (100 g, 5 min), and resuspended in BLCa organoid medium (previously described). Cells were seeded into Revvity 384-well TC-treated ViewPlate (Revvity, 6007480) in 20 µL of BLCa organoid medium at a density of 2,000 cells per well. After seeding, organoids were cultured for 7-10 days at 37°C, 5% CO₂ to facilitate organoid formation before drug treatment. Belinostat and cisplatin (SelleckChem, S1166) were freshly prepared as a 12-point, 2-fold serial dilution (10× stock) starting at 10 µM, with 20 µL of drug solution added per well (total volume 40 µL). After 72 hours of treatment, cell viability was assessed using CellTiter-Glo 3D (Promega, G9682) according to the manufacturer’s protocol with minor modifications. Plates and reagents were equilibrated to room temperature (∼30 minutes). 40 µL of CellTiter-Glo 3D reagent was added per well, followed by shaking at 375 rpm for 5 minutes, then rested for 15 minutes before luminescence was measured using a microplate reader (BioTek Synergy HTX Multimode Reader). IC₅₀ values were calculated and visualized using GraphPad Prism (version 10.4.1).

### Preparation of FFPE Tissue

Fresh PDX bladder tumor tissues were fixed in 10% neutral buffered formalin (Fisher Scientific, NC9075838) at 4°C overnight. Tissues were then dehydrated, infiltrated with wax, and embedded in paraffin blocks. FFPE samples were cut into 4-μm tissue sections and heated to 40°C on a heat plate overnight before staining.

### Immunofluorescence Staining of FFPE Sections

The sections were deparaffinized by incubating them in Citrus Clearing Solvent (Fisher Scientific, 22050122), followed by rehydration in 100%, 95%, 80%, and 70% ethanol. The slides were rinsed under running H_2_O for 2 minutes. Antigen retrieval was conducted by placing the slides in a plastic rack with pH 6.0 citrate buffer and microwaving for 20 minutes. After cooling to room temperature, the sections were outlined with a PAP pen. Dako’s blocking reagent (DAKO, S2003) was applied for 10 minutes to inhibit endogenous peroxidase, then rinsed with running H_2_O, followed by a PBS wash. The sections were blocked for one hour in a solution comprising 10% normal goat serum, 5% BSA, 0.3% Tween, and PBS. The primary antibody ARID1A (Thermo Fisher, MA5-24658) was diluted at 1:70 in the aforementioned blocking solution, applied to the sections, and incubated at room temperature overnight. The slides were washed in 1x PBST five times for 10 minutes each on the following day. Subsequently, an HRP-conjugated secondary antibody (anti-mouse, DAKO, K4001) was applied for one hour at room temperature. After three washes of 10 minutes each in PBST, the sections were treated with DAB solution (1 mL DAB substrate buffer, DAKO, K3468 + 1 drop of liquid DAB chromogen (DAKO, K4011)). The sections were rinsed with running H_2_O for 2 minutes, counterstained in hematoxylin for 90 seconds, and blued in PBS (pH 8) for 1 minute. After a final wash in running H_2_O, all sections were dehydrated in 95% and 100% ethanol and CCS before being mounted for viewing.

### The RESOLVE Trial

The RESOLVE study (NCT05154994) was a prospective, investigator-initiated phase I open-label dose-escalation clinical trial approved by the University of Utah IRB (00143952). All participants provided written informed consent. The RESOLVE study evaluated the triplet combination of tremelimumab, durvalumab, and belinostat (Figure S7). Patients with *ARID1A^m^*locally advanced or metastatic urothelial carcinoma were enrolled in a phase 1 study and planned to receive a priming dose of tremelimumab plus durvalumab (T300+D), followed by belinostat plus durvalumab for 6 cycles, then durvalumab maintenance for a total of 2 years of treatment.

## Statistical analyses

Demographic variables were analyzed using the Wilcoxon rank-sum test for age, ordinal logistic regression for stage, and the chi-square test for categorical variables. Tumor mutational burden (TMB) between patient groups was compared using Siegel regression stratified by age and sex. Time-to-event variables (overall survival and progression-free survival) were analyzed using the log-rank test. Overall survival (OS) was measured from the initial urothelial carcinoma diagnosis to death or the last contact, whichever came first. Progression-free survival (PFS) by treatment type was calculated from the start of treatment to the earliest of noted progression (including imaging tumor growth), medication change due to progression, metastasis, recurrence, or last contact, whichever occurred first. Loss to follow-up was treated as censoring.

Mutual exclusivity and co-occurrence among deleterious somatic variants were analyzed using the DISCOVER program. The significance of each gene set was determined by meta-analyzing the gene-wise p-values with Stouffer’s method. Gene sets were obtained from the MSigDB database. In total, 16197 genes and 17446 gene sets were analyzed.

R version 4.3.2 was used for statistical analysis.

## Data availability

The TCGA Bladder Urothelial Carcinoma cohort (TCGA-BLCA) WES tumor data are available from cBioPortal via the “Bladder Cancer (TCGA, Cell 2017)” dataset. The ORIEN data used in this research were generated by Aster Insights (www.asterinsights.com) in collaboration with the Oncology Research Information Exchange Network (ORIEN, www.oriencancer. org). Requests for access to the data used in this study can be submitted to the corresponding author and.

## RESULTS

### SWI/SNF alterations define a prevalent molecular subclass in urothelial carcinoma

To define the landscape of SWI/SNF (BAF) alterations in urothelial carcinoma, we analyzed whole-exome sequencing data from 792 patients in the ORIEN cohort. Deleterious alterations in SWI/SNF subunit genes (Table S1) were identified in 419 tumors (52.9%) (Figure 1A), establishing these alterations as a dominant molecular feature of urothelial carcinoma. *ARID1A* represented the most frequently altered subunit and was among the top three mutated driver genes, alongside *TP53* and *KMT2D* (Figure 1B). BAF-altered (BAF^m^) tumors were stratified into *ARID1A*-mutant (*ARID1A*^m^), non-*ARID1A* BAF-altered (OtherBAF^m^), and tumors harboring both alterations. The most mutated non-*ARID1A* subunits were *SMARCA4*, *PBRM1, ARID1B, SMARCA2,* and *ARID2*. Among all the BAF subunit variants identified, 60% were truncating variants (splice site, nonsense, or frame-shift insertion/deletion), while the others were deleterious non-truncating variants (missense or in-frame insertion/deletion). This proportion of truncating variants was significantly higher (p<0.0001) than that in the non-BAF variants (47%), with an odds ratio of 1.67 (95% confidence interval between 1.43 and 1.96). High prevalence of BAF alterations was also observed in the independent TCGA-BLCA cohort, in which 48% of tumors harbored BAF alterations, 22% had *ARID1A* alterations, and 33% had other BAF alterations (Figure S1). Collectively, these data establish that SWI/SNF alterations occur in approximately half of urothelial carcinoma tumors across independent datasets.

**Figure 1:**
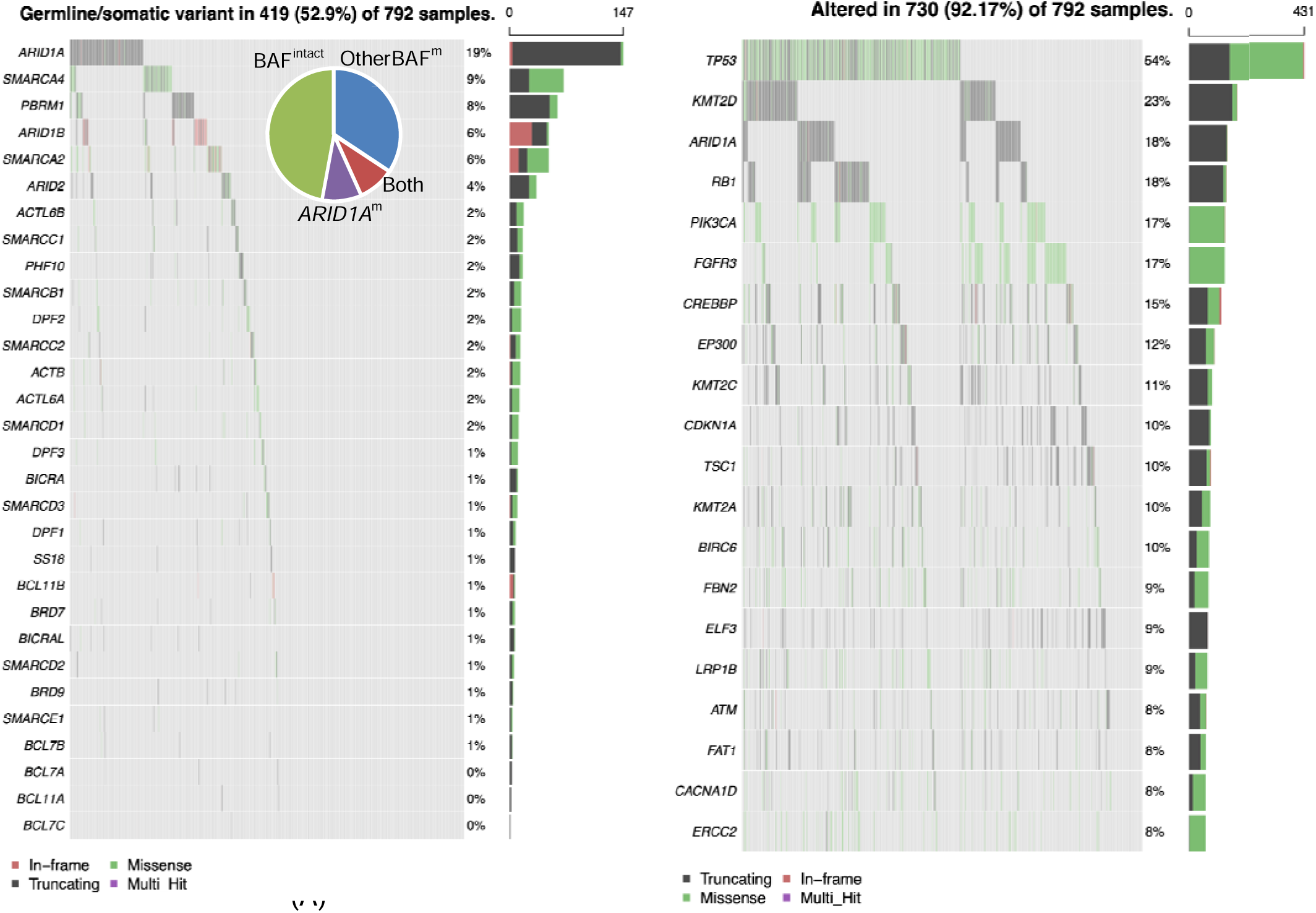
Landscape of SWI/SNF (BAF) complex alterations in urothelial carcinoma. (A) Oncoplot demonstrating deleterious germline and somatic alterations in SWI/SNF complex genes across 792 ORIEN urothelial carcinoma tumors. Alterations are colored according to mutation type. The frequency of alterations in each gene is shown on the right. (B) Oncoplot of recurrent cancer driver gene alterations in the same cohort. Only one alteration per gene per tumor was retained using the hierarchy truncating > in-frame > missense.

There were no significant differences among patients with *ARID1A*^m^, OtherBAF^m^, and those with no BAF alterations (BAF^intact^) tumors in age at diagnosis, sex, primary site, disease histology, or initial stage (Table 1). Patients with BAFO tumors showed a trend toward improved overall prognosis compared with BAF^intact^ tumors, although this difference did not reach statistical significance (Figure S2).

**Table 1:** Demographic and clinical features of patients with urothelial carcinoma in the ORIEN cohort.

|  | BAF<sup>intact</sup><br>n=373 | ARID1A<sup>m</sup><br>n=76 | OtherBAF<sup>m</sup><br>n=272 | Both<br>n=71 | P-value |
| --- | --- | --- | --- | --- | --- |
| Median age in years at diagnosis (IQR) | 68.9<br>(62.3-76.0) | 68.4<br>(62.5-76.7) | 68.9<br>(60.1-75.7) | 69.8<br>(60.3-77.3) | 0.75 |
| Male sex | 272 (72.9%) | 63 (82.9%) | 205 (75.4%) | 57 (80.3%) | 0.22 |
| Site of primary |  |  |  |  |  |
| Bladder | 345 (92.5%) | 74 (97.4%) | 250 (91.9%) | 66 (93.0%) | 0.28 |
| Upper tract (renal pelvis, ureter, kidney) | 22 (5.90%) | 1 (1.32%) | 16 (5.88%) | 4 (5.64%) |  |
| Urethra | 2 (0.54%) | 0 (0.00%) | 5 (1.84%) | 0 (0.00%) |  |
| Urachus | 3 (0.80%) | 1 (1.32%) | 1 (0.37%) | 1 (1.41%) |  |
| NOS | 1 (0.27%) | 0 (0.00%) | 0 (0.00%) | 0 (0.00%) |  |
| Histology |  |  |  |  |  |
| Urothelial | 334 (90.5%) | 74 (97.4%) | 249 (91.5%) | 68 (95.8%) | 0.63 |
| Squamous | 12 (3.25%) | 0 (0.00%) | 9 (3.31%) | 2 (2.82) |  |
| Adeno | 12 (3.25%) | 1 (1.32%) | 8 (2.94%) | 1 (1.41%) |  |
| Small cell or neuroendocrine | 11 (2.98%) | 1 (1.32%) | 6 (2.21%) | 0 (0.00%) |  |
| Stage at diagnosis |  |  |  |  |  |
| Stage 0 or 1 | 70 (23.5%) | 15 (24.6%) | 54 (23.0%) | 11 (18.6%) | 0.42 |
| Stage 2 | 68 (22.8%) | 18 (29.5%) | 73 (31.1%) | 17 (28.8%) |  |
| Stage 3 or 4 | 160 (53.7%) | 28 (45.9%) | 108 (46.0%) | 31 (52.5%) |  |

Note: IQR, interquartile range. \* P-values were derived using the Kruskal-Wallis rank sum test for age and the chi-square test for other variables.

BAF^m^ tumors were associated with significantly higher TMB compared to BAF^intact^ tumors (Figure 2A). Interestingly, the elevated TMB was primarily driven by mutations in non-*ARID1A* BAF genes rather than *ARID1A* (Figure 2B), suggesting heterogeneity in the biological consequences of distinct subunit mutations. BAF^m^ tumors in TCGA-BLCA had a significantly higher TMB too (Wilcoxon p-value <0.0001) (Figure 2C), with greater TMB associated with non-*ARID1A* mutations in the BAF complex (Figure 2D).

**Figure 2.**
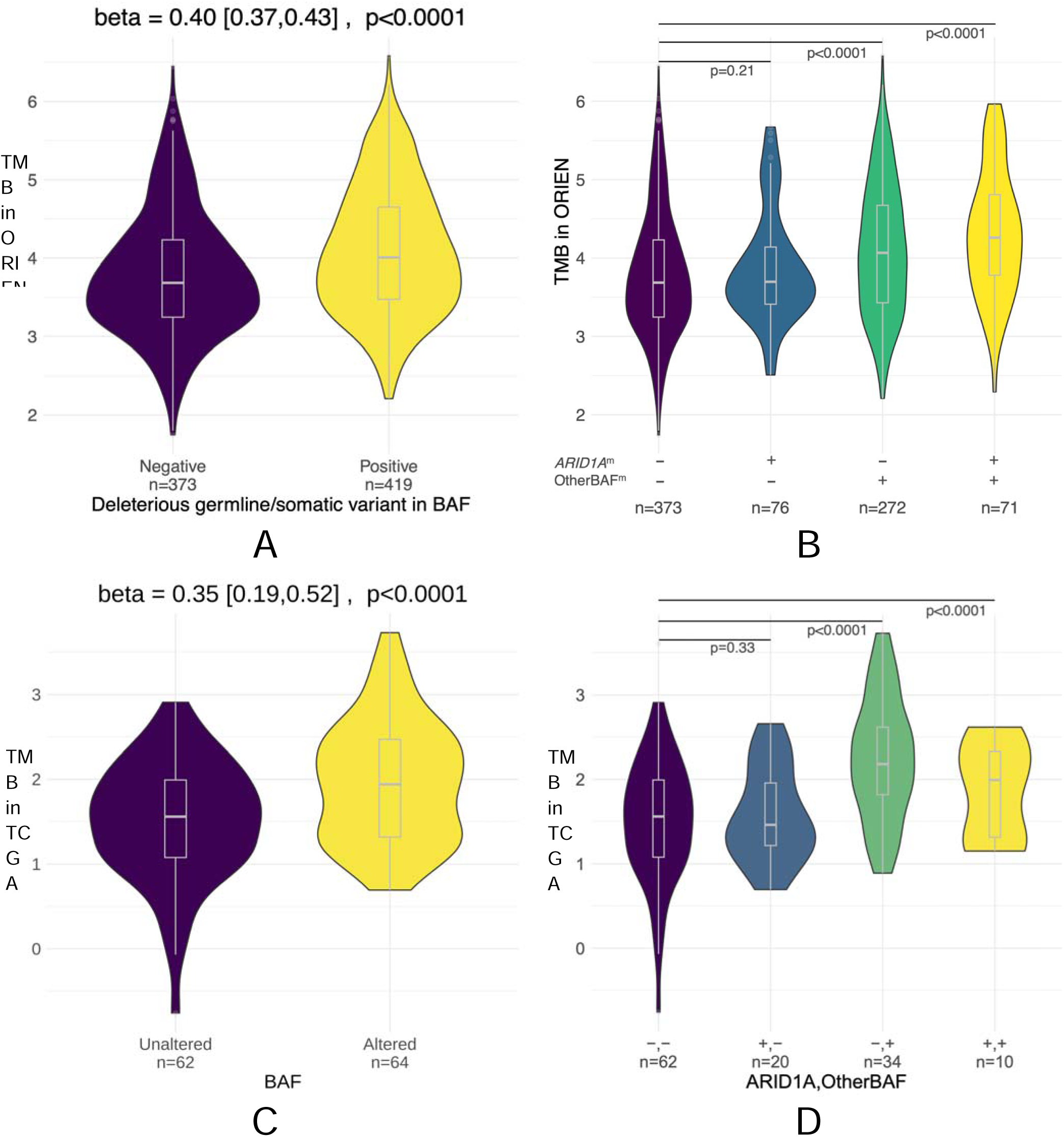
BAF-altered urothelial carcinomas exhibit increased tumor mutational burden. (A) Tumor mutational burden (TMB) in the ORIEN urothelial carcinoma cohort stratified by BAF status. Tumors harboring deleterious alterations in SWI/SNF (BAF) complex genes (BAF^m^) were compared with BAF^intact^ tumors. TMB was calculated from whole-exome sequencing data and displayed as violin plots with embedded boxplots indicating the median and interquartile range. (B) TMB in the ORIEN cohort stratified into BAF^intact^, *ARID1A*^m^, OtherBAF^m^, and tumors harboring both *ARID1A* and OtherBAF alterations. Comparisons demonstrate that increased TMB is primarily associated with non-*ARID1A* BAF alterations. (C) Validation of TMB enrichment in BAF^m^ tumors in the TCGA-BLCA cohort. (D) TMB in the TCGA-BLCA cohort stratified by BAF alteration subtype as in panel B. Whole-exome sequencing data were analyzed from 792 ORIEN tumors and 126 TCGA-BLCA tumors. Beta coefficients and corresponding *P* values were derived using Siegel regression adjusted for age and sex. Boxplots depict median and interquartile range; violin plots represent sample density distributions. For panel B, group sizes were n=373 (BAF^intact^), n=76 (*ARID1A*^m^), n=272 (OtherBAF^m^), and n=71 (Both). For panel D, group sizes were n=62 (BAF^intact^), n=20 (*ARID1A*^m^), n=34 (OtherBAF^m^), and n=10 (Both). Horizontal bars indicate pairwise comparisons. Exact *P* values are shown in the figure.

No significant difference in co-occurring mutations was identified between BAF^m^ and BAF^intact^ tumors. We observed nominal mutual exclusivity (raw p < 0.05) in 58 genes (Table S2) and several gene sets (Table S3). However, none of these associations remained statistically significant after correction for multiple testing (Benjamini–Hochberg FDR > 0.05). The nominally associated genes and pathways, which include several previously reported in urothelial carcinoma (e.g., *FGFR3, KMT2D, KDM6A*), are presented in Tables S2 and S3.

### BAF alterations converge on a reproducible transcriptional program characterized by proliferation and loss of differentiation

To determine whether BAF alterations define a biologically distinct state, we first performed transcriptomic analyses using RNA sequencing data from the ORIEN cohort and validated the findings in TCGA-BLCA. Compared with BAF^intact^ tumors, BAF^m^ tumors showed enrichment for proliferation-associated pathways, including E2F targets, the G2M checkpoint, and spermatogenesis (Figure S3B). In contrast, immune-related pathways (TNF-α signaling via NF-κB, TGF-β signaling) and cellular stress responses (hypoxia and reactive oxygen species pathways) were downregulated (Figure S3B).

We next examined whether these patterns differed by mutation subtype. *ARID1A*^m^ tumors showed enrichment of immune signaling, particularly interferon-γ responses (Figure S3C). In contrast, OtherBAF^m^ were characterized by broad depletion of immune activation pathways, including interferon-α and interferon-γ responses, allograft rejection, TNF-α signaling via NF-κB, and inflammatory response (Figure S3D). Notably, cell cycle activation (E2F targets) was consistently enriched across both *ARID1A*^m^ and OtherBAF^m^ tumors (Figure S3C, S3D).

These findings were validated in an independent TCGA-BLCA cohort using a parallel approach (Figure S4), and meta-analysis across both datasets confirmed the robustness of the transcriptional program (Figure 3). Proliferation-related pathways (E2F targets and the G2M checkpoint) were uniformly upregulated across cohorts (Figure 3B), reinforcing the notion that BAF alterations drive cell-cycle activation and chromatin dysregulation in urothelial carcinoma. BAF^m^ tumors consistently demonstrated downregulation of metabolic pathways, particularly the reactive oxygen species pathway, relative to BAF^intact^ tumors (Figure 3B). This suggests a shared metabolic vulnerability that is consistent across cohorts, regardless of which subunit is mutated.

**Figure 3.**
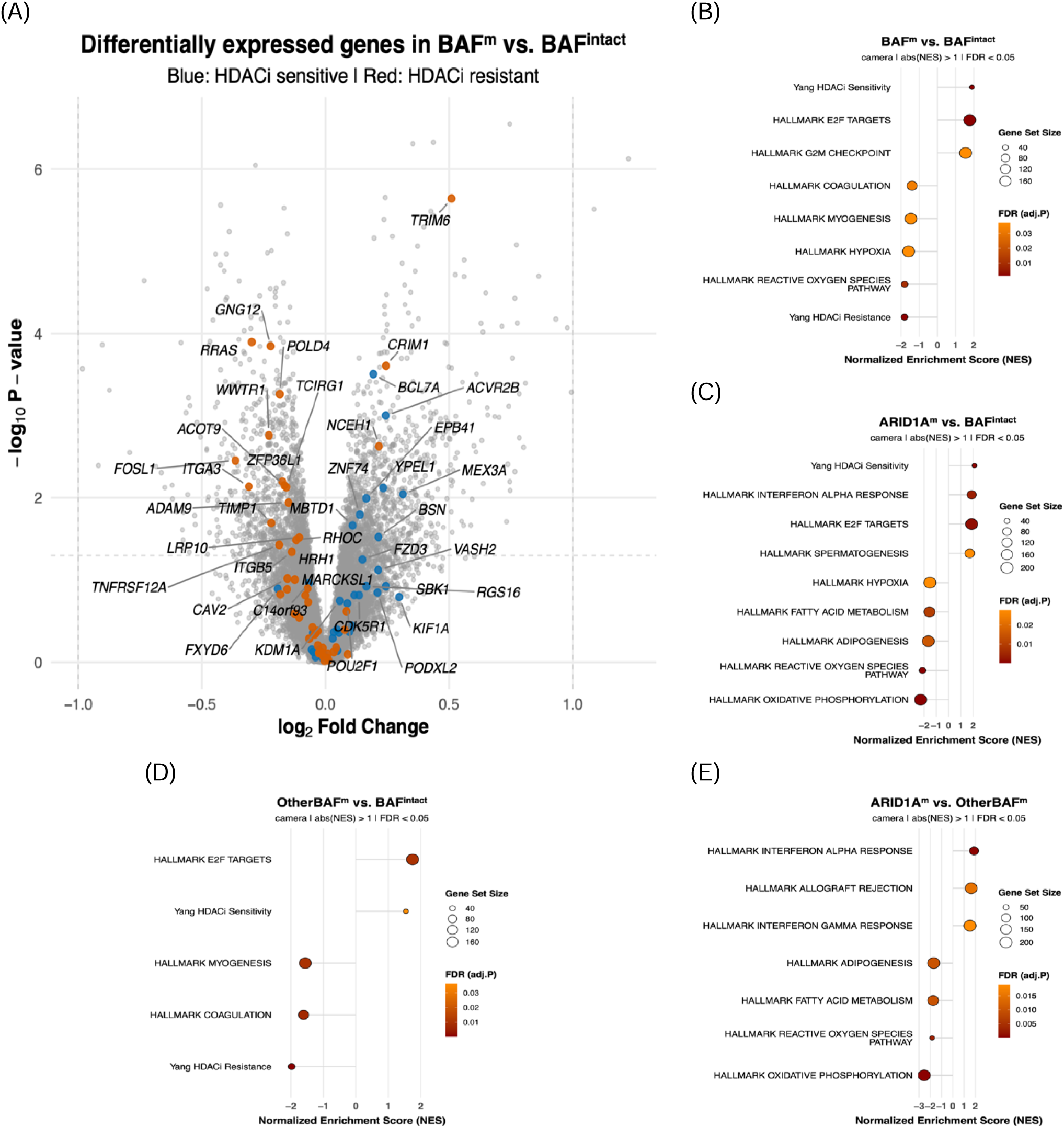
BAF alterations define a convergent transcriptional state associated with proliferati n, loss of differentiation programs, altered metabolic signaling, and predicted sensitivity to histone deacetylase (HDAC) inhibition. (A) Meta-analysis volcano plot integrating ORIEN and TCGA-BLCA transcriptomic datasets comparing BAF^m^ and BAF^intact^ tumors. Genes belonging to previously described HDAC inhibitor sensitivity and resistance signatures are highlighted. Blue points denote HDAC inhibitor-sensitive signature genes and orange points denote HDAC inhibitor-resistance signature genes. (B) Meta-analysis gene-set enrichment analysis (GSEA) comparing BAF^m^ with BAF^intact^ tumors. Positive normalized enrichment scores (NES) indicate enrichment in BAF^m^ tumors, whereas negative scores indicate depletion. Hallmark pathways and HDAC inhibitor sensitivity/resistance signatures are shown. (C) GSEA comparing *ARID1A*^m^ tumors with BAF^intact^ tumors. (D) GSEA comparing OtherBAF^m^ tumors with BAF^intact^ tumors. (E) GSEA directly comparing *ARID1A*^m^ and OtherBAF^m^ tumors. Transcriptomic analyses were performed using RNA sequencing data from the ORIEN cohort and validated in the TCGA-BLCA cohort. Differential expression analyses were conducted using limma, and pathway enrichment analyses were performed using CAMERA. Meta-analysis across cohorts was performed using Stouffer’s method. Bubble size corresponds to gene-set size, and color intensity indicates false-discovery rate (FDR-adjusted *P* value). Only pathways meeting significance thresholds of |NES| >1 and FDR <0.05 are displayed.

In addition, consistent depletion of the myogenesis gene set in BAF^m^ tumors (Figure 3B) indicates loss of differentiation programs and increased cellular plasticity. To define the specific molecular drivers underlying the observed depletion of the myogenesis pathway, we examined the meta-analysis results for individual genes in this signature (Figure S5). A strong downregulation was observed for *WWTR1* (encoding the transcriptional co-activator TAZ), a central effector of the YAP/TAZ signaling axis. This was accompanied by consistent loss of several key structural and regulatory components, including the actin-associated scaffolding protein *PDLIM7*, the myosin light chain *MYL6B*, the tropomyosin *TPM2*, and the transcription factor *SPDEF*. Additional significantly depleted transcripts included *TGFB1, BIN1, AK1,* and *SPEG*, further supporting impaired cytoskeletal organization and myogenic differentiation programs in BAF^m^ tumors.

Finally, to directly compare transcriptional differences between subtypes, we performed differential expression analysis between *ARID1A*^m^ and OtherBAF^m^ tumors. *ARID1A*^m^ tumors were enriched for immune activation, whereas OtherBAF^m^ tumors demonstrated enrichment of metabolic pathways (Figure 3E), supporting the presence of biologically distinct subgroups within BAF^m^ disease.

### BAF^m^ tumors exhibit transcriptional signatures predictive of HDAC inhibitor sensitivity

Given the central role of SWI/SNF complexes in chromatin regulation, we hypothesized that BAF-altered tumors may exhibit therapeutic vulnerability to chromatin modulation. To test this, we interrogated established pan-cancer transcriptomic signatures associated with sensitivity and resistance to histone deacetylase (HDAC) inhibition.^14^

BAF^m^ tumors demonstrated significant enrichment for the HDAC inhibitor-sensitive signature and a concurrent depletion of the HDAC inhibitor-resistant signature compared to BAF-intact controls (Figure 3A, 3B). These enrichments and depletions of signatures were consistent across the ORIEN and TCGA cohorts (Figures S4A, S4B, S5A, and S5B). Importantly, HDAC inhibitor sensitivity was observed in both *ARID1A^m^* and OtherBAF^m^ tumors (Figures 3C, 3D), suggesting that HDAC inhibitor sensitivity may be a conserved vulnerability across different BAF complex mutations associated with disruption of the SWI/SNF complex rather than a mutation-specific effect.

At the gene level, depletion of the HDAC inhibitor-resistance signature in BAF^m^ tumors was primarily driven by coordinated downregulation of multiple interconnected genes, including the YAP/TAZ pathway component *WWTR1* (TAZ), key regulators of cell adhesion and migration (*ITGB5, ITGA3, ADAM9, CAV2, RHOC, RRAS,* and *GNG12*), pro-inflammatory cytokine signaling components (*TNFRSF12A*), and the AP-1 transcription factor subunit *FOSL1* (Figure 3A).

### *NECTIN4* expression and molecular subtype of BAF^m^ urothelial carcinoma

Beyond the hyper-proliferation of BAF^m^ tumors, we identified additional molecular features with therapeutic relevance. Specifically, analysis of the ORIEN cohort revealed that BAF^m^ tumors exhibit slightly higher *NECTIN4* expression than BAF^intact^ controls (Figure 4A). A consistent, significant trend was also observed in the TCGA cohort (Figure 4B). Molecular subtypes of the ORIEN urothelial carcinoma tumors were categorized by the consensus muscle-invasive bladder cancer (cMIBC) classifier. The luminal class was more frequently observed among BAF^m^ (46%) than BAF^intact^ (35%) tumors (Figure 4C, 4D).

**Figure 4.**
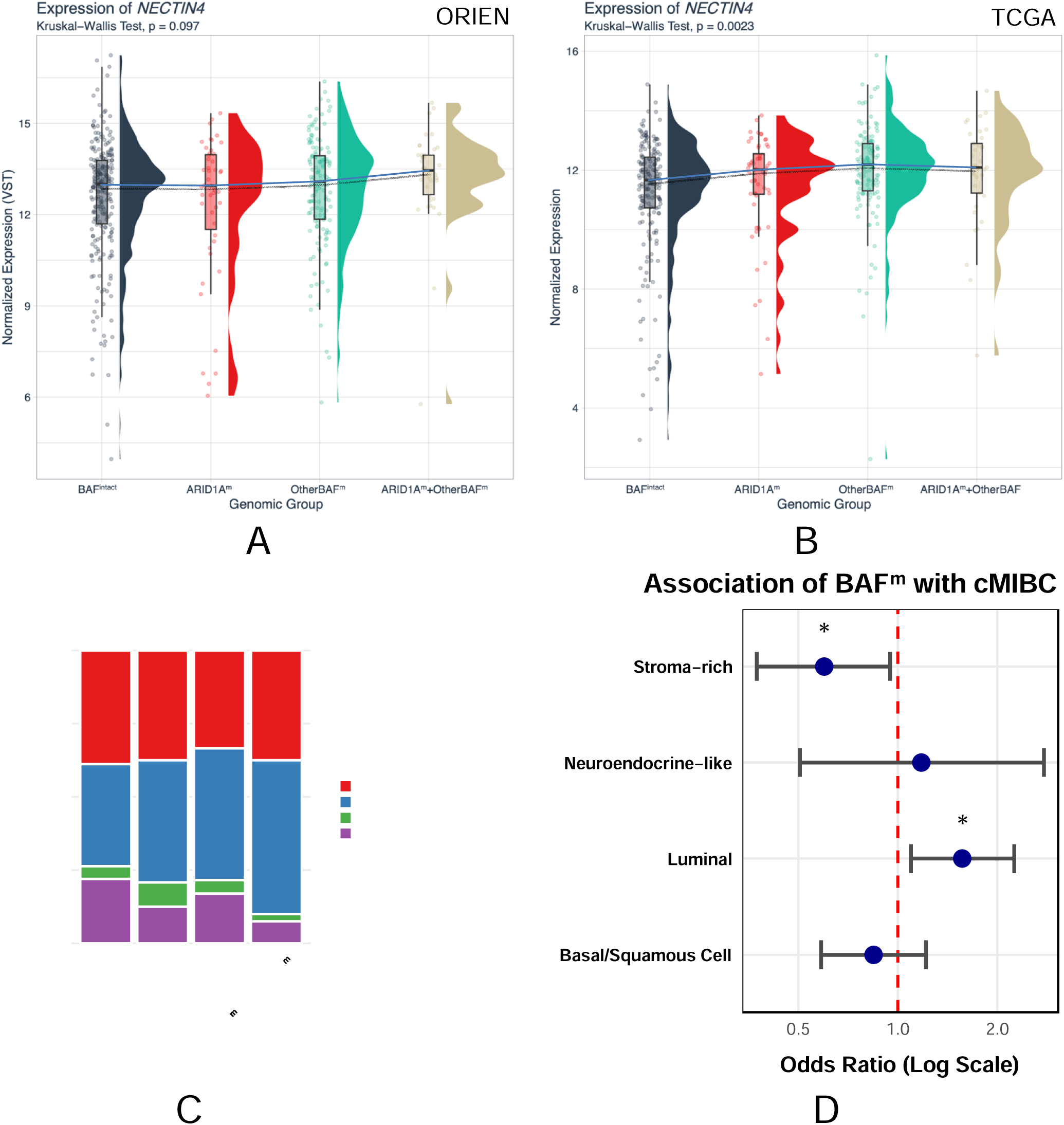
BAF alterations are associated with elevated NECTIN4 expression and enrichment for luminal molecular subtypes. (A) *NECTIN4* mRNA expression in the ORIEN urothelial carcinoma cohort stratified by BAF alteration status. (B) Validation of *NECTIN4* expression patterns in the TCGA-BLCA cohort. (C) Distribution of consensus muscle-invasive bladder cancer (cMIBC) molecular subtypes accordin to BAF status in the ORIEN cohort. (D) Association between BAF alterations and cMIBC molecular classification. RNA sequencing data were obtained from ORIEN and TCGA-BLCA tumors. *NECTIN4* expression values are shown as variance-stabilized normalized counts. Boxplots depict median and interquartile range, while violin plots indicate expression density distributions. Individual points represent tumors. Group comparisons were performed using the Kruskal–Wallis test, with exact *P* values indicated in the figure. BAF^m^ tumors demonstrated enrichment for luminal molecular subtype features and increased *NECTIN4* transcript expression relative to BAF^intact^ tumors. cMIBC, consensus molecular classification of muscle-invasive bladder cancer.

### HDAC inhibition remodels chromatin accessibility and suppresses oncogenic transcriptional programs

Given that *ARID1A* is the most frequently altered BAF subunit and has the strongest existing translational evidence for HDAC inhibitor sensitivity, subsequent mechanistic studies were performed in *ARID1A*^m^ models. To define the mechanistic basis of HDAC inhibitor sensitivity, we performed integrated transcriptomic and chromatin accessibility profiling following treatment with the pan-HDAC inhibitor belinostat in *ARID1A*^m^ urothelial carcinoma cell line HT1197.

RNA sequencing revealed widespread transcriptional reprogramming characterized by downregulation of key oncogenic pathways, including E2F, MYC, G2M checkpoint, epithelial–mesenchymal transition, and DNA repair programs (Figure 5A). These changes directly mirror the proliferative transcriptional programs observed in BAF^m^ tumors, supporting concordance between tumor state and therapeutic vulnerability (Figure 3B).

**Figure 5:**
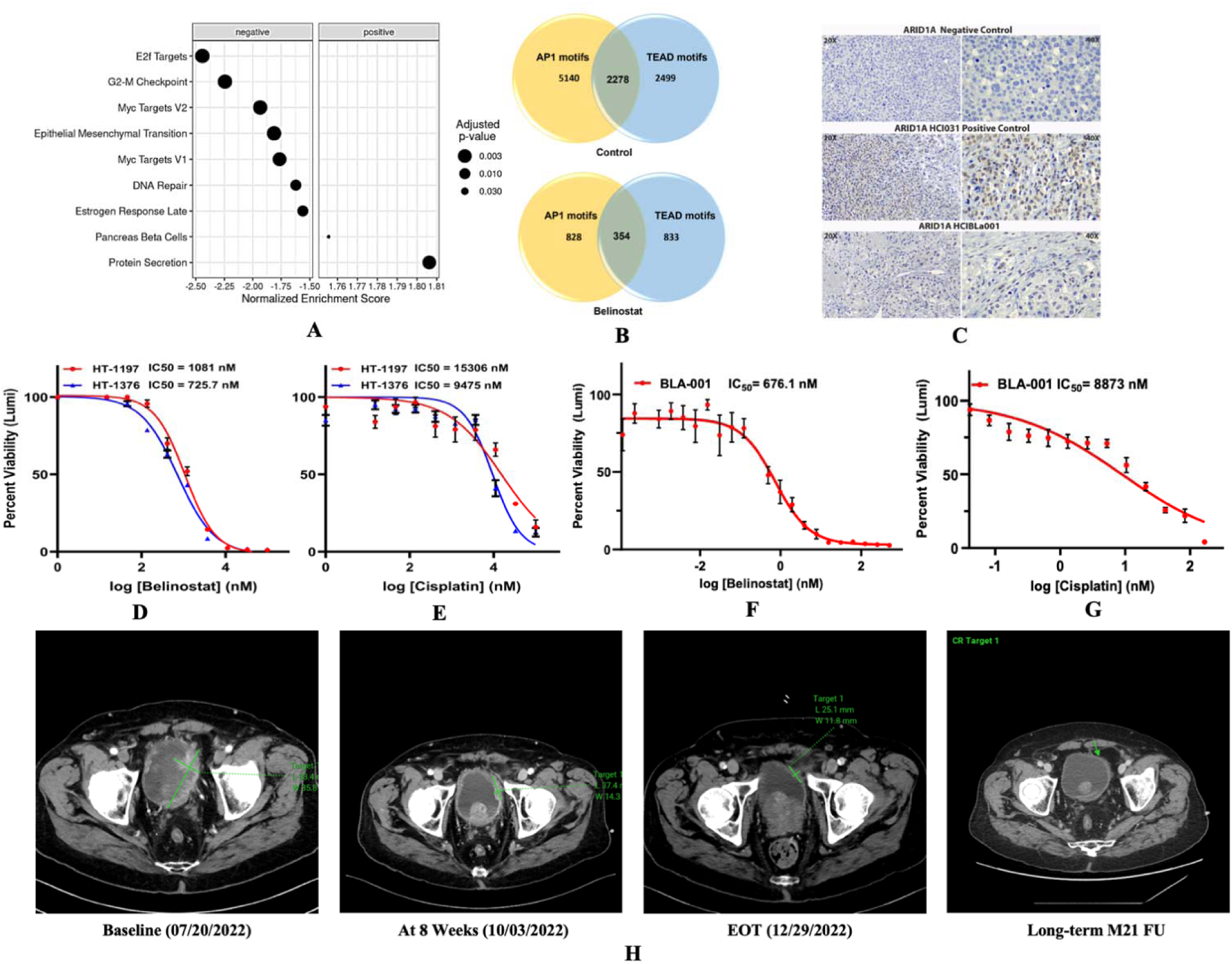
HDAC inhibition disrupts AP-1/TEAD-driven transcriptional programs and demonstrates therapeutic activity in ARID1A-mutant urothelial carcinoma. (A) Gene-set enrichment analysis of RNA-seq data from belinostat-treated HT-1197 cells demonstrating suppression of E2F targets, G2M checkpoint, MYC targets, epithelial–mesenchymal transition, DNA repair, and additional oncogenic transcriptional programs. (B) ATAC-seq motif analysis of HT-1197 cells demonstrating overlap between accessible chromatin regions containing AP-1 and TEAD transcription factor motifs in untreated and belinostat-treated conditions. Belinostat treatment reduced the number of accessible AP-1/TEAD-containing regions. (C) Representative ARID1A immunohistochemistry demonstrating loss of ARID1A protein expression in the corresponding patient-derived xenograft tumor. Positive and negative controls are shown for comparison. (D,E) Dose–response curves demonstrating the effects of belinostat and cisplatin on viability of *ARID1A*^m^ urothelial carcinoma cell lines HT-1197 and HT-1376. (F,G) Dose–response analyses of belinostat and cisplatin in *ARID1A*^m^ PDxOs generated from a cisplatin-resistant urothelial carcinoma. (H) Representative serial imaging from a patient with *ARID1A*^m^ urothelial carcinoma treated on the RESOLVE trial with tremelimumab, durvalumab, and belinostat, demonstrating durable disease control. For cell viability studies, HT-1197 and HT-1376 cells were treated with serial dilutions of drug for 72 hours and viability was measured using CellTiter-Glo. PDxO viability was assessed using CellTiter-Glo 3D following 72 hours of treatment. Experiments were performed in triplicate (n=3 biological replicates per condition), and values are presented as mean ± SEM. RNA-seq and ATAC-seq analyses were performed in three independent biological replicates per condition. Differential gene expression was analyzed using DESeq2, and motif enrichment was performed using HOMER. The radiographic response shown in panel H represents an individual patient enrolled on the RESOLVE clinical trial and illustrates preliminary clinical translation of the observed preclinical findings.

To determine whether these transcriptional changes were accompanied by remodeling of chromatin structure, we performed ATAC-seq. Genome-wide profiling identified 79,286 merged accessible regions across conditions, with a net reduction in chromatin accessibility following belinostat treatment. Specifically, 4,825 regions demonstrated increased accessibility, whereas 14,618 regions showed decreased accessibility relative to untreated controls (Figure S6A), indicating global chromatin closure.

Integration of chromatin accessibility with gene expression data demonstrated concordance between chromatin and transcriptional changes. Regions gaining accessibility were associated with transcriptional activation (n=2,100), while regions losing accessibility correlated with gene downregulation (n=3,000) (Figure S6), supporting a model in which belinostat-induced transcriptional reprogramming is driven, in part, by direct modulation of chromatin accessibility at target loci.

To define the regulatory networks underlying these effects, we performed motif enrichment analysis across all accessible chromatin regions. At baseline, accessible chromatin was strongly enriched for transcription factor motifs associated with proliferation and oncogenic signaling, including AP-1 family members (FOSL2, FRA1, FRA2, JUNB) and TEAD transcription factors (TEAD1, TEAD3), as well as architectural and lineage regulators such as CTCF and RUNX1/2. Notably, belinostat treatment was associated with enrichment of alternative transcriptional programs, including motifs for CHOP, FOXK1, FOXL2, and OCT2.

Given the prominent baseline enrichment of AP-1 and TEAD motifs, we next examined whether belinostat functionally disrupted these pathways by assessing motif-associated chromatin accessibility. Among 79,286 accessible regions, 2,278 AP-1/TEAD motif-containing sites were accessible in untreated cells; however, this number decreased markedly to 354 following belinostat treatment (Figure 5B). This selective loss of accessibility indicates targeted repression of AP-1- and TEAD-regulated enhancers and promoters.

Consistent with the known roles of AP-1 and TEAD in mediating proliferation, survival, and immune evasion through MEK/ERK and YAP/TAZ signaling pathways, their selective chromatin closure is associated with downregulation of cell-cycle, proliferation, and epithelial–mesenchymal transition programs observed in transcriptomic analyses.

Collectively, these findings demonstrate that HDAC inhibition remodels the chromatin landscape in *ARID1A*^m^ urothelial carcinoma through preferential repression of AP-1/TEAD-driven regulatory networks, thereby suppressing key transcriptional dependencies that sustain the *ARID1A*^m^ tumor state.

### Functional validation confirms sensitivity to HDAC inhibition in *ARID1A*^m^ preclinical models

Building on observed HDAC inhibitor vulnerability signatures and prior clinical observations that patients with *ARID1A*^m^ urothelial carcinoma^15^ we evaluated sensitivity to histone deacetylase inhibitor belinostat in *ARID1A*^m^ urothelial carcinoma cell lines. To extend these findings to a more clinically relevant system, we generated PDxOs from an *ARID1A*^m^ urothelial carcinoma tumor that had demonstrated resistance to neoadjuvant cisplatin-based chemotherapy. Loss of ARID1A protein expression in the corresponding PDX tumor was confirmed by immunohistochemistry (Figure 5C).

In 2D cell line models (HT-1197 and HT-1376), belinostat induced a dose-dependent reduction in cell viability. Notably, belinostat demonstrated substantially greater potency than cisplatin, with mean IC₅₀ values of 1081.9 nM and 725.7 nM, compared with 15,306 nM and 9,475 nM for cisplatin, respectively (Figures 5D,E). These data indicate enhanced sensitivity to HDAC inhibition relative to standard cytotoxic therapy.

Consistent with cell line data, belinostat treatment significantly reduced PDxO viability, with an IC₅₀ of 540.9 nM (Figure 5F), whereas cisplatin exhibited minimal activity, with an IC₅₀ of 57,150 nM (Figure 5G), consistent with the resistant phenotype.

Collectively, these results demonstrate that sensitivity to HDAC inhibition is maintained in clinically relevant, chemotherapy-resistant models of *ARID1A*^m^ urothelial carcinoma, supporting the potential therapeutic utility of HDAC inhibitors in treatment-refractory disease.

### RESOLVE clinical trial to treat *ARID1A*^m^ urothelial carcinoma with a triplet of tremelimumab, durvalumab, and belinostat

To explore clinical translation, we analyzed data from the investigator-initiated RESOLVE trial (NCT05154994; Figure S7), evaluating a combination regimen, a triplet regimen comprising tremelimumab (CTLA-4 inhibitor), durvalumab (PD-L1 inhibitor), and belinostat (HDAC inhibitor) in patients with advanced urothelial carcinoma harboring *ARID1A*^m^. Among the initial cohort of 3 enrolled patients (July 2022 and January 2023), one patient achieved a durable radiographic response with prolonged disease control lasting more than 2 years (Figure 5H). However, subsequent enrollment was limited by immune-mediated toxicity associated with dual checkpoint blockade in extensively pretreated patients with *ARID1A*^m^ urothelial carcinoma, leading to modification of the trial design.

## DISCUSSION

Our study of 1200 urothelial carcinoma tumors across two independent datasets demonstrates that disruption of the SWI/SNF (BAF) chromatin remodeling complex defines a distinct molecular subclass of urothelial carcinoma. This subclass of BAF^m^ urothelial carcinoma is characterized by a reproducible genomic state. Although alterations occur across multiple BAF subunits (Figure 1A), these diverse mutations converge on a shared program marked by increased TMB (Figure 2), plus a transcriptional signature characterized by activation of E2F-, G2M-, and MYC- driven proliferation, loss of lineage identity, and dysregulation of metabolic pathways (Figure 3). Prior studies have focused primarily on *ARID1A*, which has been shown to broadly impair chromatin remodeling and to be associated with higher TMB.^16^ Our findings establish BAF alterations as state-defining genomic events that create a unified, therapeutically targetable tumor context in approximately half of urothelial carcinoma cases. Importantly, this chromatin-disrupted state is consistently associated with transcriptomic signatures predictive of sensitivity to HDAC inhibition (Figure 3), supporting a model in which chromatin dependencies represent a central and actionable vulnerability in BAF^m^ urothelial carcinoma.

A central insight from this study is that diverse alterations across BAF subunits converge on a shared transcriptional phenotype, rather than producing subunit-specific effects. While *ARID1A* represents the most frequently mutated component of the complex, our analyses indicate that impairment of chromatin remodeling capacity itself is the dominant determinant of tumor state. This reframes BAF alterations as drivers of a common chromatin-disrupted program characterized by transcriptional plasticity and altered chromatin-dependent gene regulation.

Recent studies have demonstrated that *ARID1A* loss is associated with upregulation of cell cycle and DNA repair programs in bladder cancer organoid models, supporting a role for SWI/SNF dysfunction in driving proliferative transcriptional states.^17^ Our findings are consistent with these observations and extend them by demonstrating that this transcriptional phenotype reflects a broader, chromatin-defined dependency shared across diverse BAF subunit alterations, rather than an *ARID1A*-specific phenomenon.

Mechanistically, our integrated transcriptomic and chromatin accessibility analyses provide insight into how this chromatin-disrupted state can be therapeutically targeted. HDAC inhibition induced widespread remodeling of chromatin accessibility, characterized by a net reduction in accessible regions and preferential suppression of AP-1 and TEAD transcription factor binding sites (Figure 5B). These regulatory networks are key mediators of proliferation, survival, and cellular plasticity through MEK/ERK and YAP/TAZ signaling pathways. Their selective suppression was associated with coordinated downregulation of E2F, MYC-, and epithelial-mesenchymal transition programs (Figure 5A), establishing a direct link between chromatin remodeling and transcriptional control of tumor behavior. These findings align with prior studies demonstrating that the SWI/SNF complex functions as a regulator of YAP/TAZ activity and constrains oncogenic transcriptional programs at enhancer regions.^18^ In addition, genome-wide analyses have shown that YAP/TAZ-TEAD complexes co-occupy enhancers with AP-1 transcription factors to drive the expression of proliferation-associated genes.^19^ Our data extend this framework by demonstrating that HDAC inhibition selectively disrupts these AP-1/TEAD-associated chromatin regions in *ARID1A*^m^ urothelial carcinoma, thereby providing a mechanistic link among SWI/SNF dysfunction, chromatin-dependent gene regulation, and therapeutic vulnerability. These findings support a model in which HDAC inhibition targets chromatin-dependent transcriptional programs associated with SWI/SNF dysfunction, thereby disrupting key dependencies of the BAF^m^ tumor state.

From a translational perspective, these findings are directly relevant to contemporary treatment paradigms in urothelial carcinoma. Sensitivity to HDAC inhibitors was preserved across preclinical models, including patient-derived organoids generated from cisplatin-resistant tumors (Figure 5D-G), supporting the therapeutic potential of targeting chromatin dependencies in treatment-refractory disease. Prior studies have demonstrated that *ARID1A^m^* cancers are vulnerable to HDAC inhibitors, including a dependence on HDAC activity for tumor survival and response to chromatin remodelling therapies.^20,21^ Our findings build on this work by providing mechanistic evidence linking HDAC inhibitor sensitivity to disruption of AP-1/TEAD-driven chromatin accessibility and transcriptional programs.

Clinical development of HDAC inhibitors in urothelial carcinoma historically been guided by alterations in histone acetylation regulators, including histone acetyltransferase genes such as *CREBBP* and *EP300*. However, biomarker-directed approaches based on individual epigenetic modifiers have yielded limited clinical success. For example, the phase II study of the HDAC inhibitor mocetinostat in patients with advanced urothelial carcinoma harboring acetyltransferase pathway alterations demonstrated only modest activity and did not establish a clinically actionable predictive biomarker.^22^ In contrast, our findings suggest that the therapeutically relevant determinant of HDAC inhibitor sensitivity may not be disruption of individual acetylation regulators, but rather impairment of the SWI/SNF chromatin remodeling complex. Despite substantial genetic heterogeneity across BAF subunits, BAF^m^ tumors converged on a reproducible transcriptional and chromatin-defined state characterized by activation of proliferative programs, loss of differentiation programs, and enrichment of HDAC inhibitor sensitivity signatures. Importantly, a recent clinical study evaluating HDAC inhibitor-based combination with PD-1 blockade (tucidinostat plus tislelizumab) in platinum resistant advanced urothelial carcinoma reported objective response rates approaching 50%,^23^ rekindling interest in chromatin-targeting strategies in this disease, although predictive biomarkers were not reported. Collectively, these observations raise the possibility that SWI/SNF alterations may represent a more biologically coherent biomarker class for patient selection than individual histone acetyltransferase gene alterations and provide a mechanistic framework for future biomarker-enriched trials of HDAC inhibitor–based combinations in urothelial carcinoma.

Notably, we observed an association between BAF alterations and *NECTIN4* expression, suggesting that this chromatin-defined tumor state may be enriched for the expression of therapeutically targetable surface antigens. NECTIN4 is the target of the antibody-drug conjugate enfortumab vedotin, and prior studies have demonstrated that NECTIN4 expression is enriched in luminal molecular subtypes of urothelial carcinoma and is a key determinant of enfortumab vedotin sensitivity.^24^ Our observation that BAF^m^ tumors are enriched for luminal features and exhibit increased *NECTIN4* expression suggests a potential biologic link between chromatin state, lineage identity, and therapeutic target expression. This correlation needs to be validated at the protein level before being interpreted for treatment prediction. While this hypothesis requires further validation, it provides a biologically grounded framework for biomarker-enriched clinical trial design within the current therapeutic landscape.

Preliminary clinical translation of this approach is supported by observations from the RESOLVE trial, in which a patient with *ARID1A*^m^ urothelial carcinoma achieved a durable response to an HDAC inhibitor in combination with immune checkpoint blockade (5H). Although limited by sample size and complicated by immune-related toxicity associated with dual checkpoint inhibition, this observation provides early clinical support for the therapeutic relevance of targeting chromatin vulnerabilities in this population. These data underscore the need for optimized combinatorial strategies that retain efficacy while minimizing toxicity, and strategic sequencing of therapies.

We note some limitations of our study. We intentionally prioritized mechanistic studies in *ARID1A*^m^ urothelial carcinoma models because *ARID1A* alterations represent the most common BAF-complex event in urothelial carcinoma and provided the strongest available translational bridge between cohort-level observations, available laboratory models, and an ongoing biomarker-enriched clinical trial. While this approach enabled detailed mechanistic investigation, it also represents a limitation, as additional studies will be required to establish whether other BAF-complex alterations exhibit identical chromatin regulatory dependencies. Furthermore, the association between BAF alterations and NECTIN4 expression was assessed at the transcript level and will require protein-level validation to establish its relevance for therapeutic targeting. Finally, prospective clinical studies are needed to determine the appropriate stage and line of therapy and to define optimal combination strategies incorporating HDAC inhibitors in biomarker-selected patient populations.

In summary, we define a convergent chromatin-dependent tumor state across BAF^m^ urothelial carcinoma characterized by proliferative signaling, loss of differentiation, and altered chromatin accessibility. This state is associated with a reproducible vulnerability to HDAC inhibitors and is supported by mechanistic evidence demonstrating selective repression of AP-1/TEAD-driven transcriptional programs. Together, these findings provide a conceptual and translational framework for targeting chromatin dependencies in urothelial carcinoma and support the development of biomarker-guided therapeutic strategies, including rational combinations with contemporary agents such as antibody-drug conjugates.

## Acknowledgments

Research reported in this article was partially supported by the U.S. Department of Veterans Affairs Merit Review Award #1T01 BX005765 (to SG). The contents do not represent the views of the U.S. Department of Veterans Affairs or the United States Government. The research reported here used the Total Cancer Care protocol, the High-Throughput Genomics Shared Resource, the Preclinical Cancer Models Shared Resource (PCM), and the Biorepository & Molecular Pathology (BMP) Shared Resource at the Huntsman Cancer Institute, University of Utah. Immunohistochemistry (IHC) studies were performed by Erica Egal and her team at the Research Immunohistochemistry Core. Research reported in this publication utilized the Cancer Bioinformatics Shared Resource at the Huntsman Cancer Institute at the University of Utah and was supported by the National Cancer Institute of the National Institutes of Health under Award Number P30CA042014. Computational resources used in this work were partially funded by the NIH Shared Instrumentation Grant 1S10OD021644-01A1. The content is solely the responsibility of the authors and does not necessarily represent the official views of the NIH. The RESOLVE clinical trial was conducted with support from the Clinical Trials Office at the Huntsman Cancer Institute and was funded by AstraZeneca and Acrotech. During the preparation of this manuscript, the authors used Microsoft copilot for language editing and clarity. The authors reviewed and edited all content and take full responsibility for the final manuscript.

## Author contributions

Conception or design of study: S.Gu.B.R.C; Acquisition of data: K.F. and S.Gu.; Analysis of data: B.-J.F., K.F., D.A.N., A.At., C.Cap., C.J.S., D.H.L., T.J.P., and C.Car.; Interpretation of data: B.-J.F., K.F., D.A.N., A.At., C.Cap., C.J.S., D.H.L., T.J.P., C.Car., D.G., L.G., E.A.S., K.N., Z.M., E.K., J.K., S.Gh., P.H., P.V.V., A.Ay., M.L.C., U.S., N.A., B.R.C., and S.Gu.; Drafting the article or revising it critically: B.-J.F., K.F., and S.Gu.; Accountability for all aspects of the work: B.-J.F., K.F., D.A.N., A.At., C.Cap., C.J.S., D.H.L., T.J.P., C.Car., D.G., L.G., E.A.S., K.N., Z.M., E.K., J.K., S.Gh., P.H., P.V.V., A.Ay., M.L.C., U.S., N.A., B.R.C., and S.Gu.

