## Supplementary Material for "SWI/SNF Alterations Define a Chromatin-Dependent Subtype of Urothelial Carcinoma"

### Supplement

Figure S1: Oncoplot of BAF mutations in TCGA-BLCA.

Figure S2: Clinical outcomes of patients with BAF^m^ vs. BAF^intact^ tumors in ORIEN.

Figure S3 A: gene set enrichment analysis comparing BAF^m^ to BAF^intact^ tumors in ORIEN.

Figure S4: Analysis of TCGA’s RNA-seq data.

Figure S5: Meta-analysis result of Hallmark Myogenesis genes.

Figure S6: Integrated transcriptomic and epigenomic analysis of belinostat-treated HT-1197 cells.

Figure S7: RESOLVE study schema.

Table S1: BAF genes.

Table S2: Genes demonstrating mutual exclusivity with BAF^m^ identified in ORIEN tumor exomes.

Table S3: Pathways demonstrating mutual exclusivity with BAF^m^ identified in ORIEN tumor exomes.

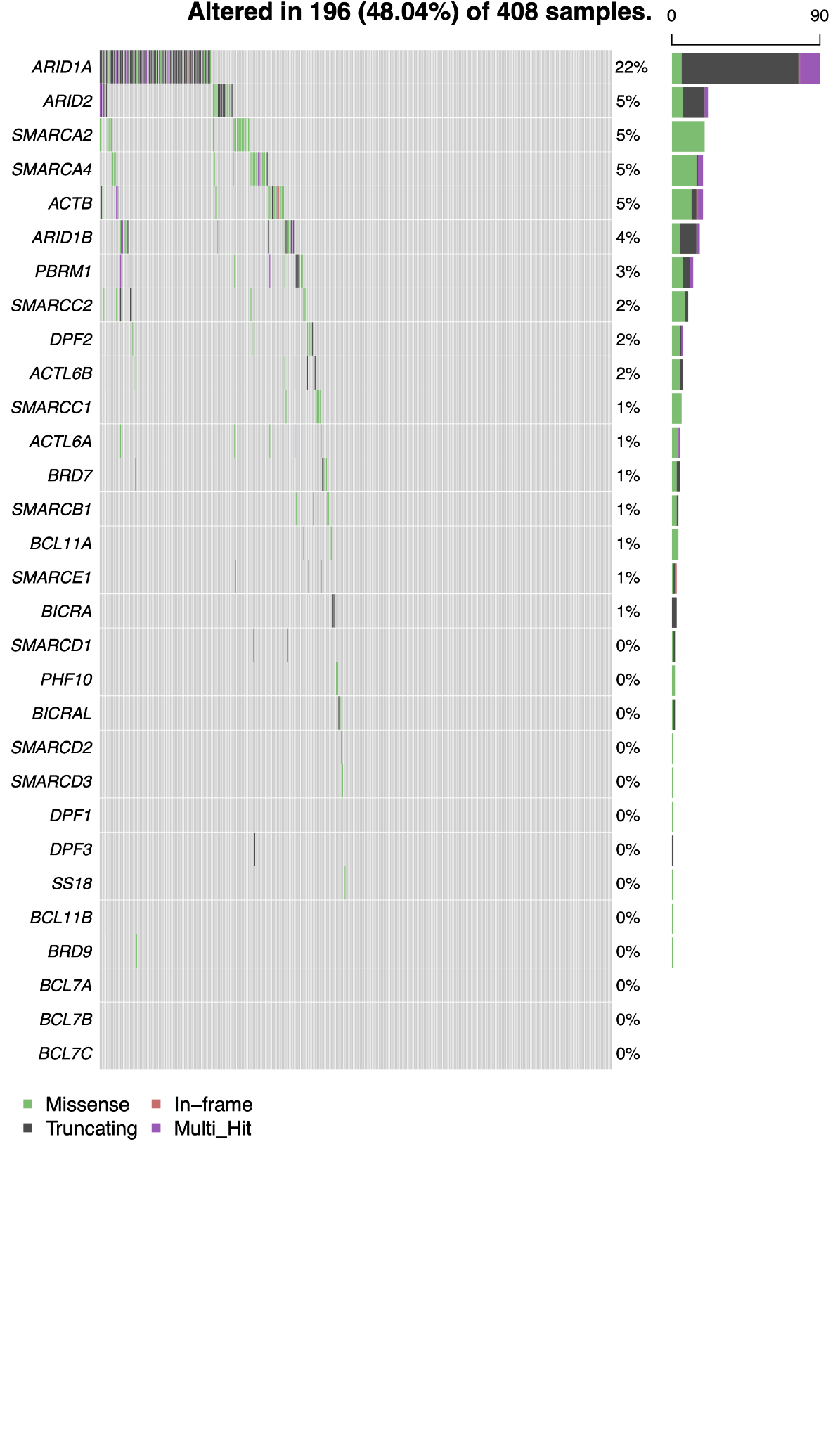

Figure S1: Oncoplot of BAF mutations in TCGA-BLCA.

| 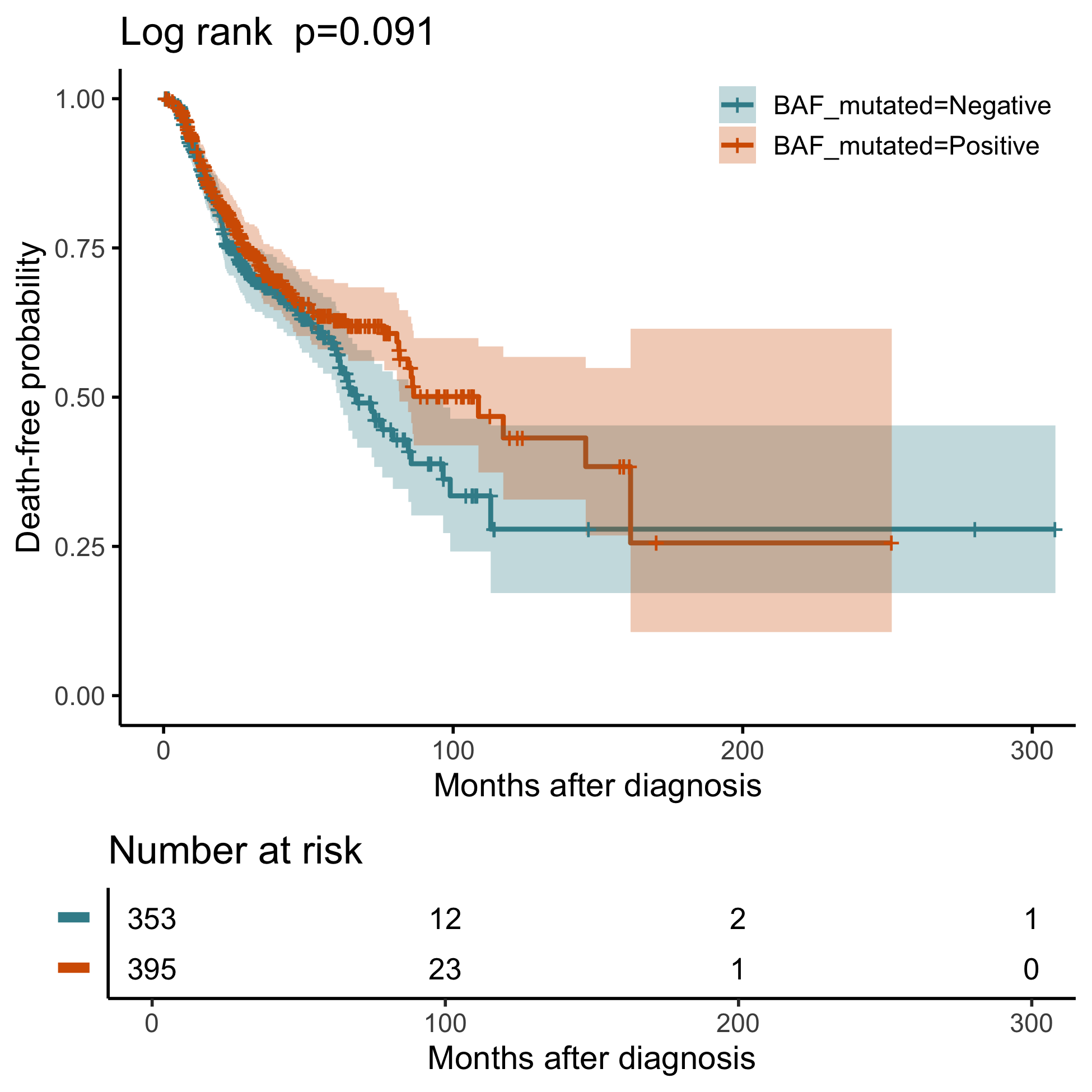 | 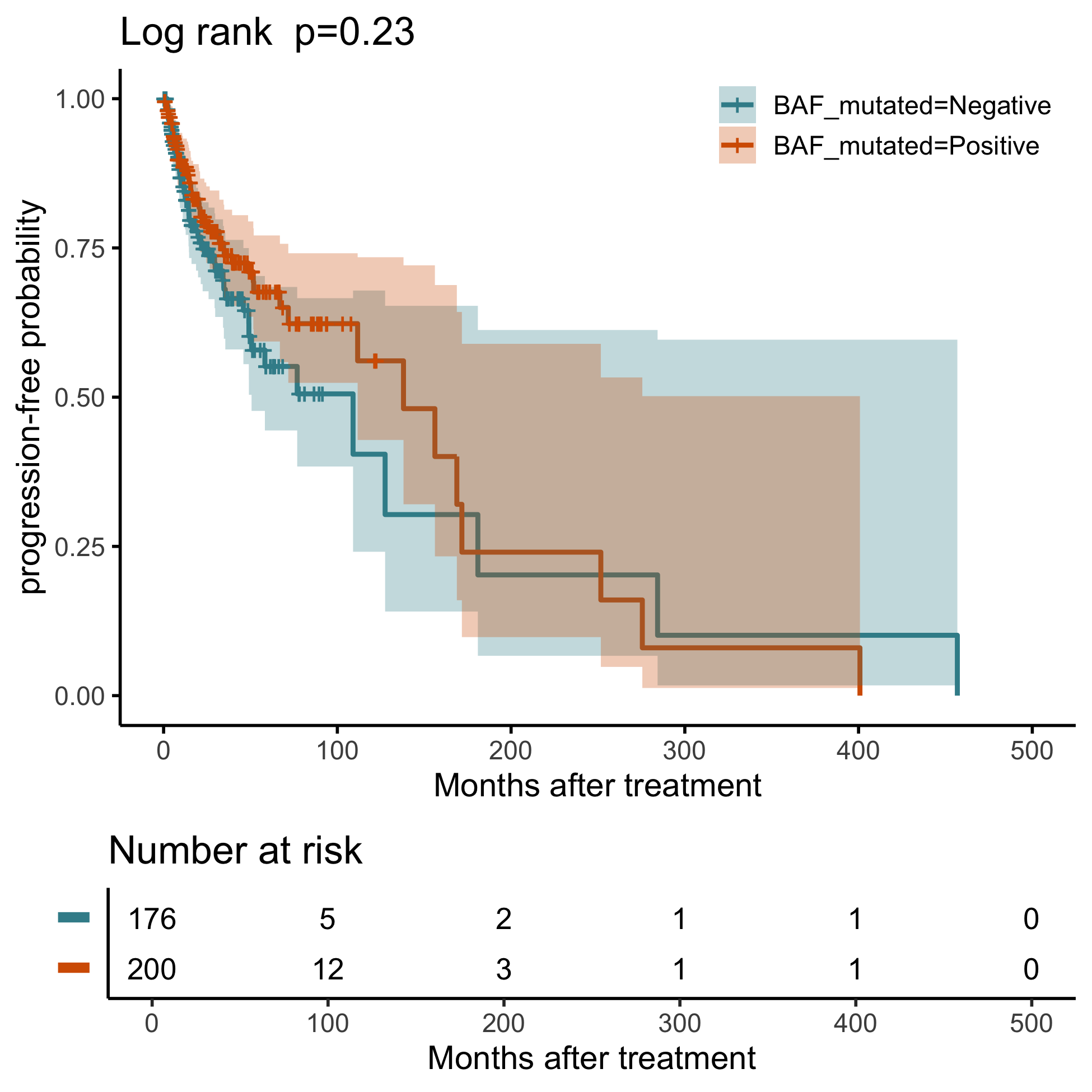 |
| --- | --- |
| A | B |
| 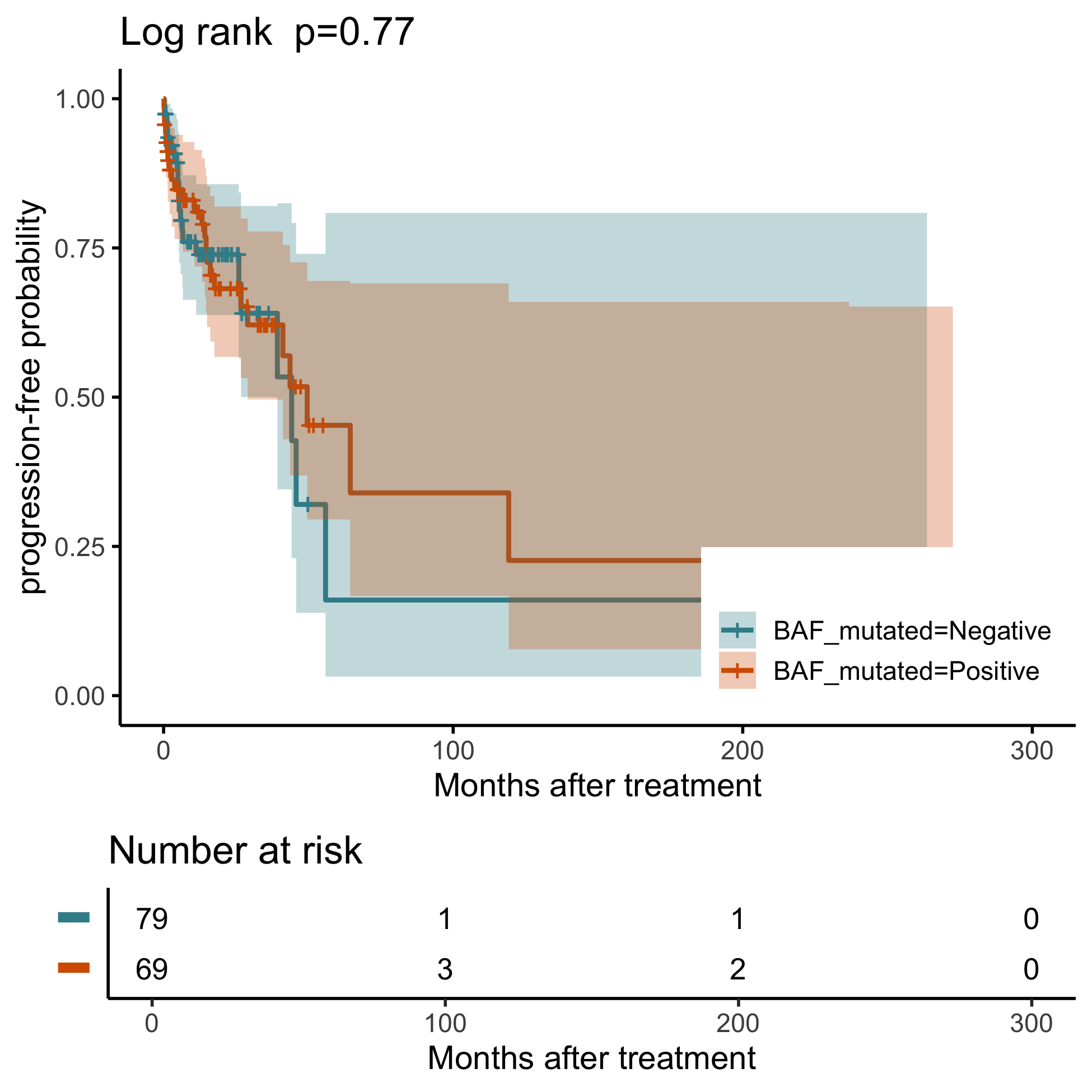 | 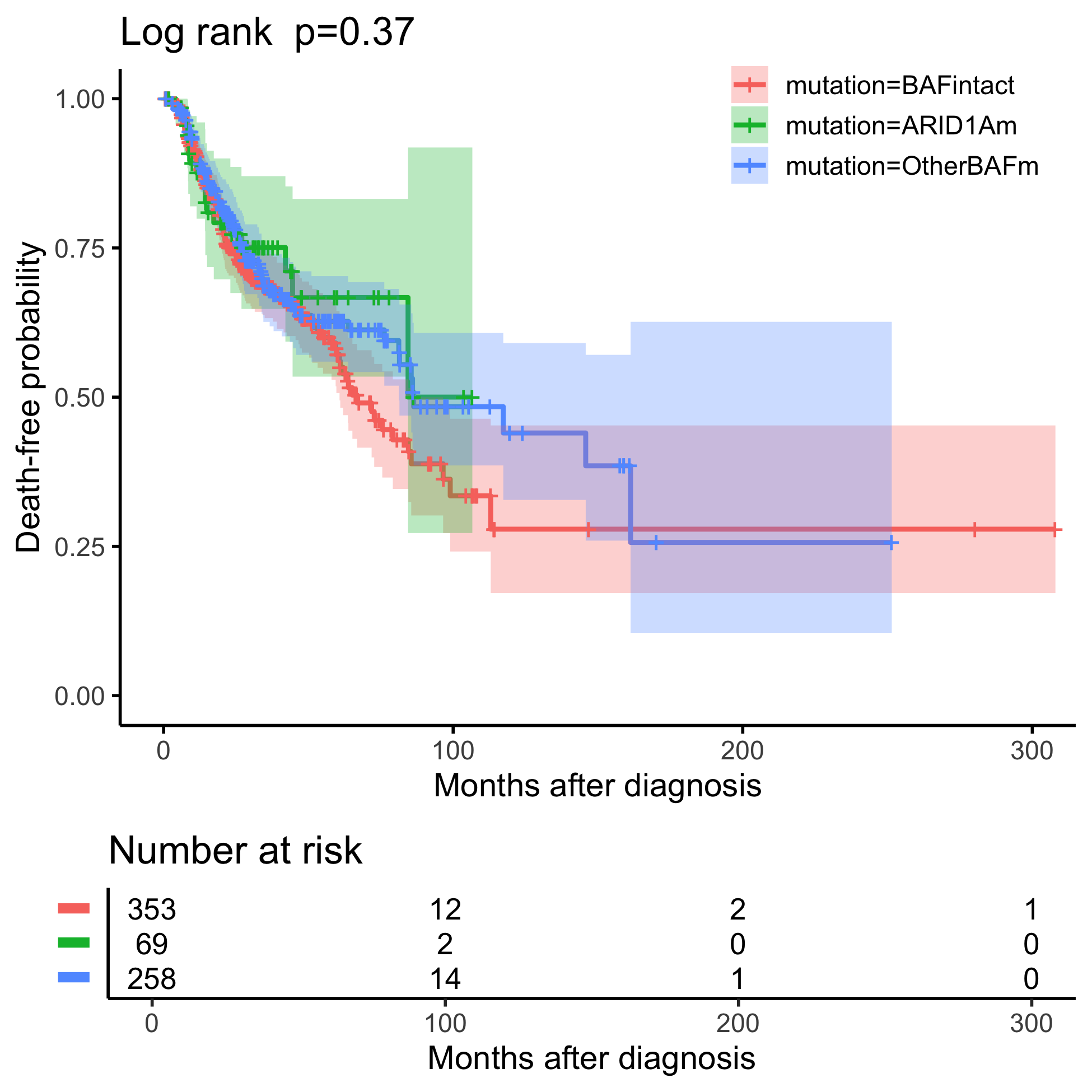 |
| C | D |

Figure S2: Clinical outcomes of patients with BAF^m^ vs. BAF^intact^ tumors in ORIEN. (A) overall survival, (B) progression-free survival after platinum-based treatment, (C) progression-free survival after ICI, (D) overall survival, separating patients into three groups.

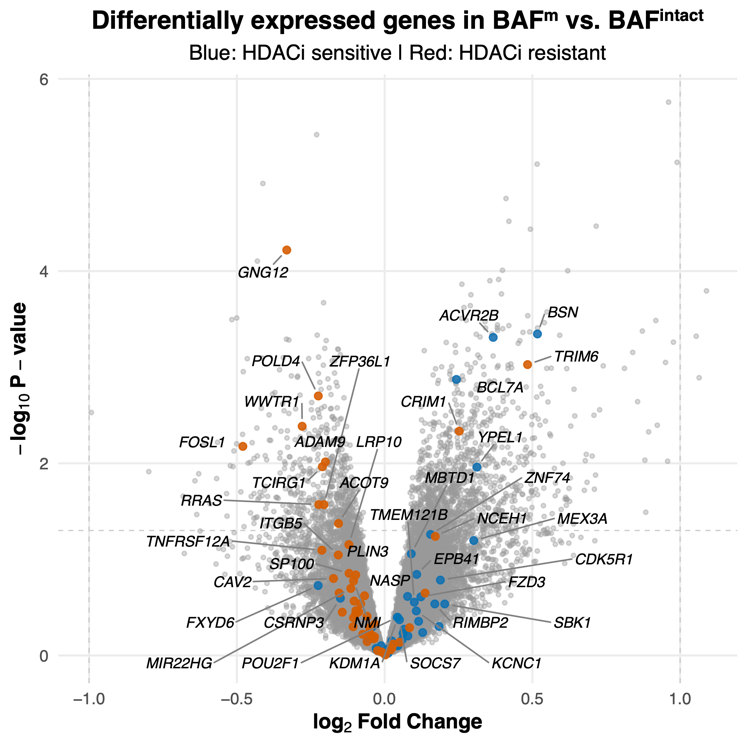

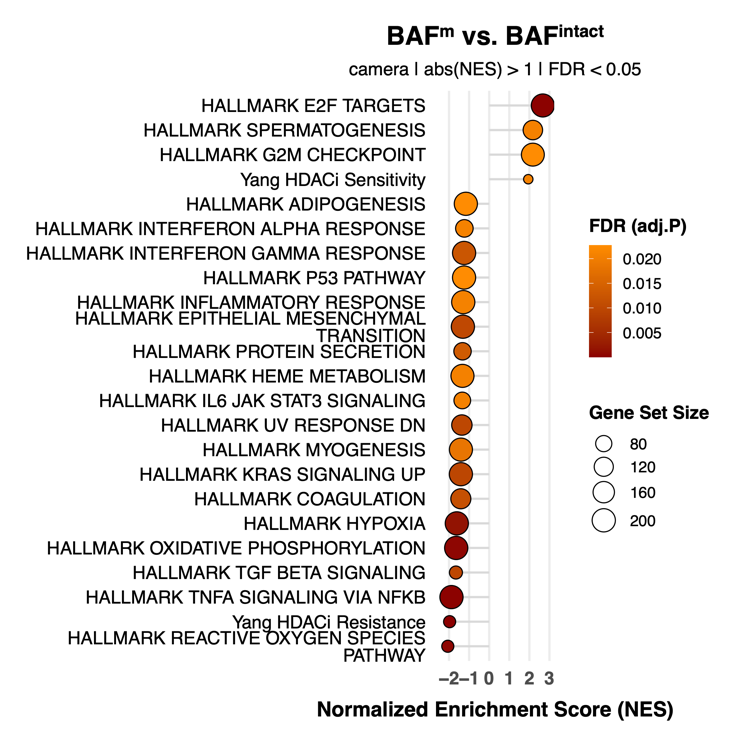

A	B

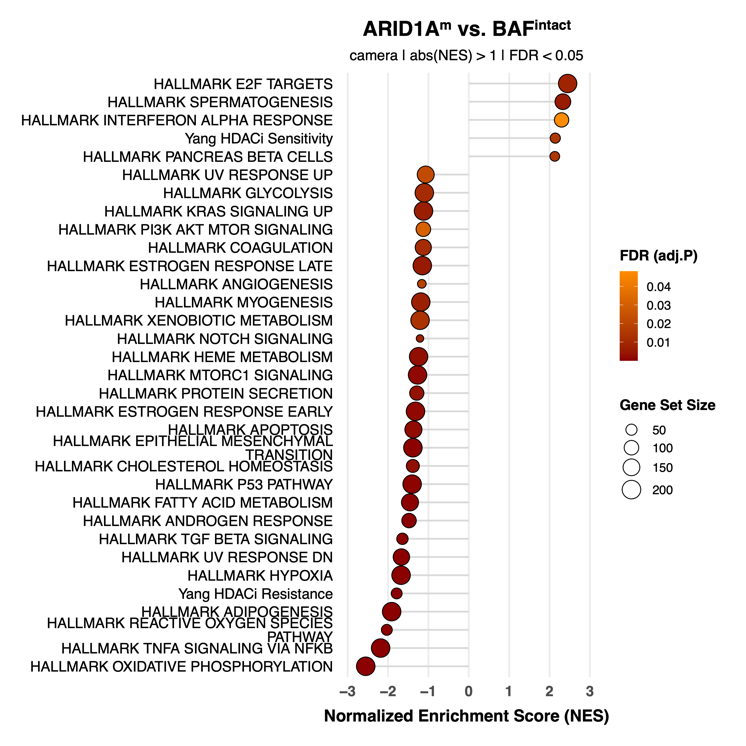

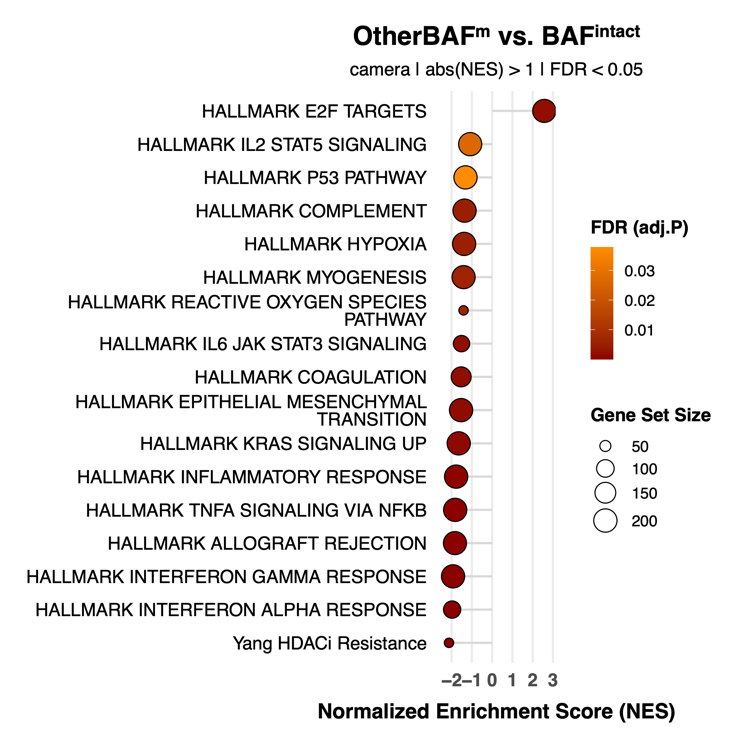

C	D

Figure S3: Analysis of ORIEN’s RNA-seq data. (A) volcano plot of each gene’s result comparing BAF^m^ to BAF^intact^ tumors; (B) gene set enrichment analysis comparing BAF^m^ to BAF^intact^ tumors; (C) gene set enrichment analysis comparing *ARID1A^m^* to BAF^intact^ tumors; (D) gene set enrichment analysis comparing OtherBAF^m^ to BAF^intact^ tumors.

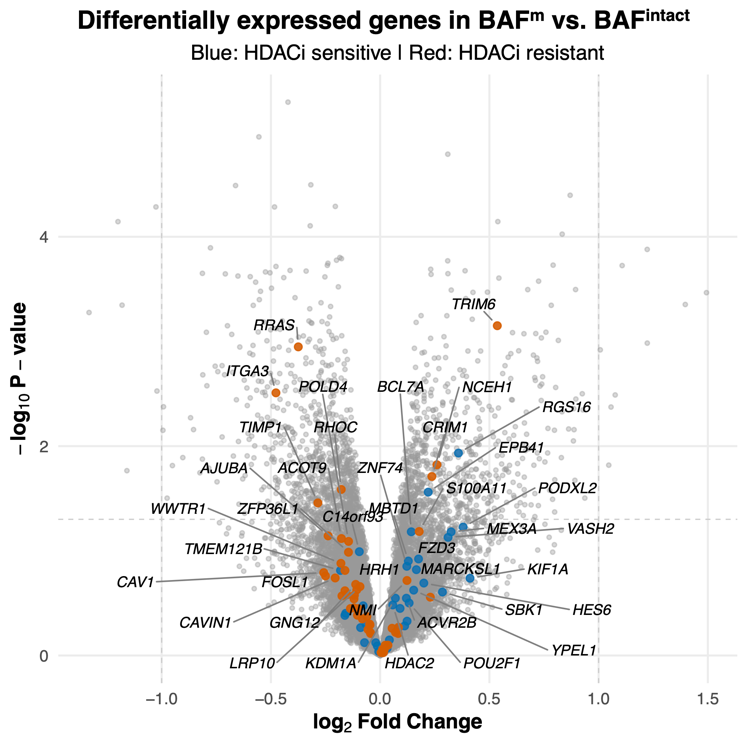

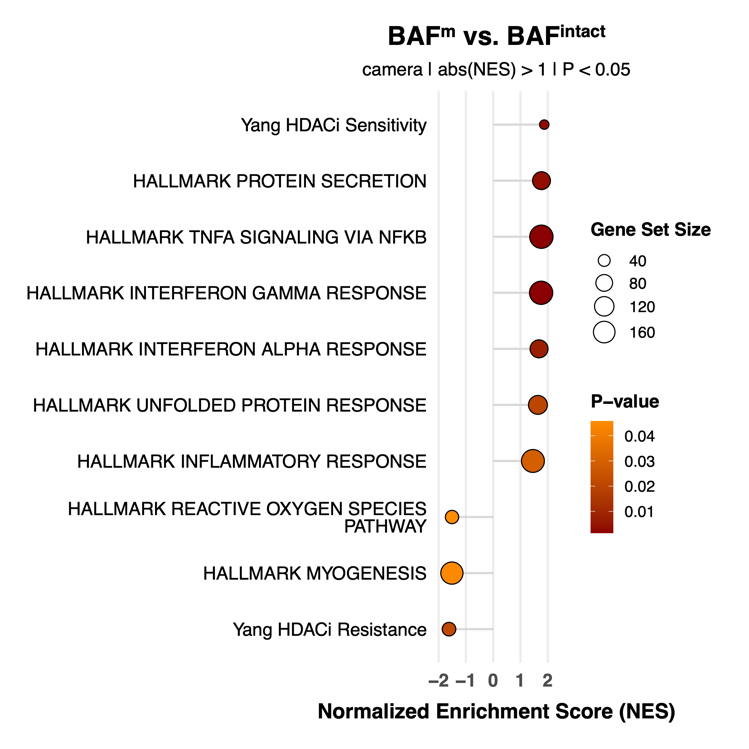

| A | B |
| --- | --- |

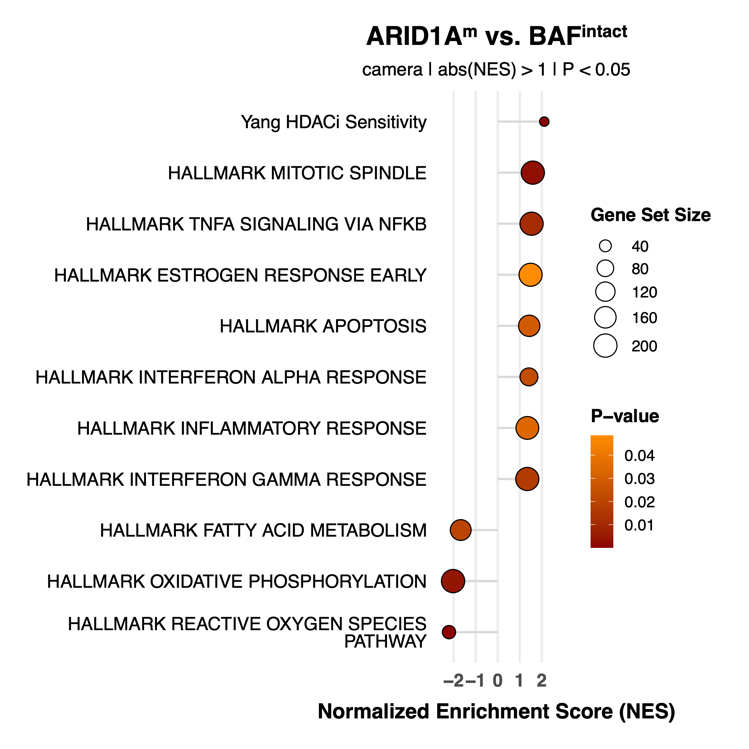

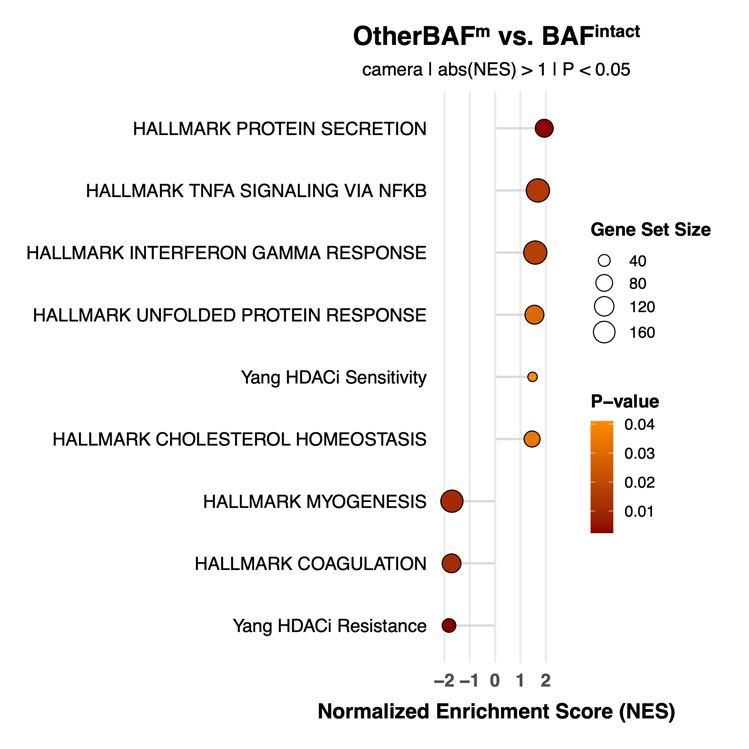

| C | D |
| --- | --- |

Figure S4: Analysis of TCGA’s RNA-seq data. (A) volcano plot of each gene’s result comparing BAF^m^ to BAF^intact^ tumors; (B) gene set enrichment analysis comparing BAF^m^ to BAF^intact^ tumors; (C) gene set enrichment analysis comparing *ARID1A^m^* to BAF^intact^ tumors; (D) gene set enrichment analysis comparing OtherBAF^m^ to BAF^intact^ tumors.

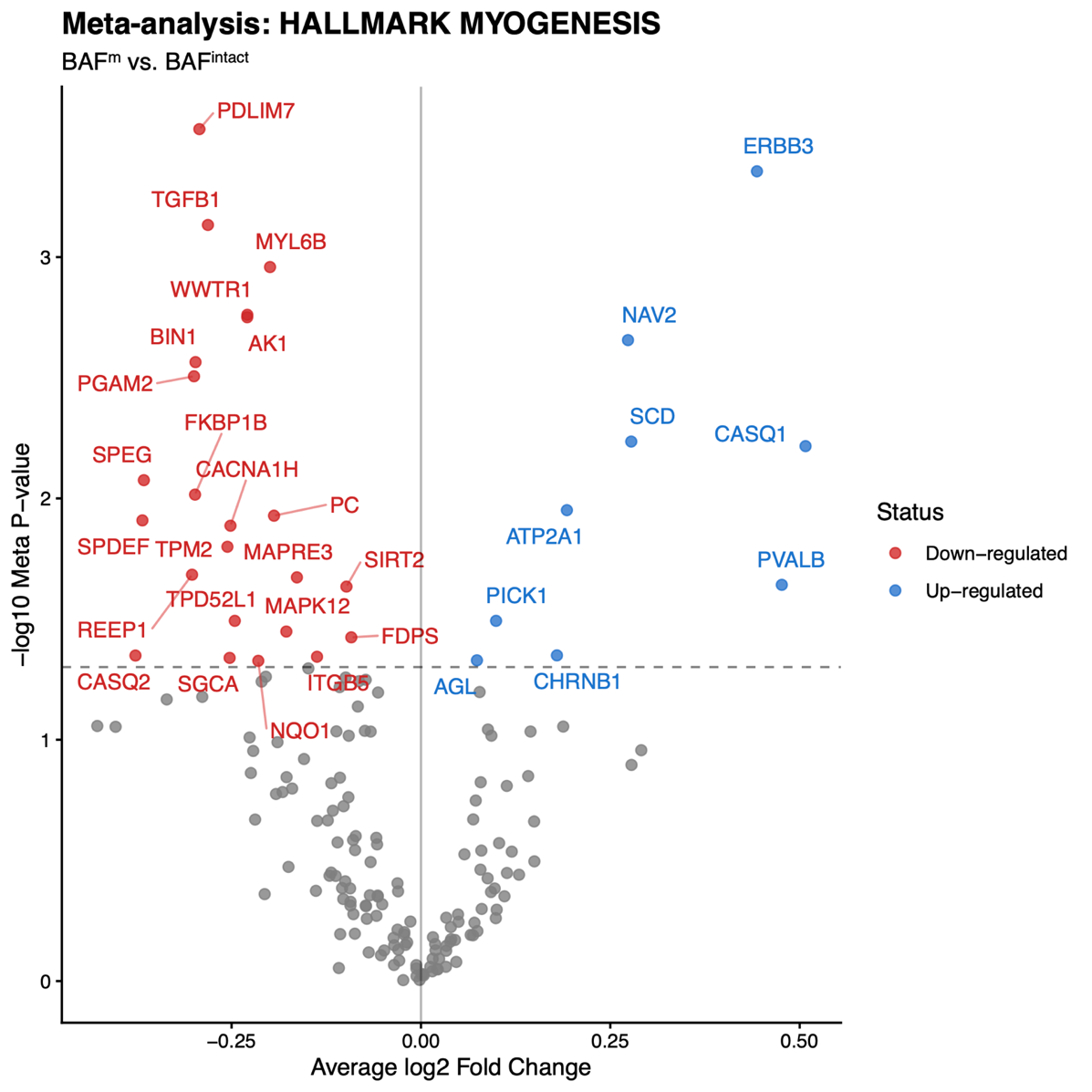

Figure S5: Meta-analysis result of Hallmark Myogenesis genes.

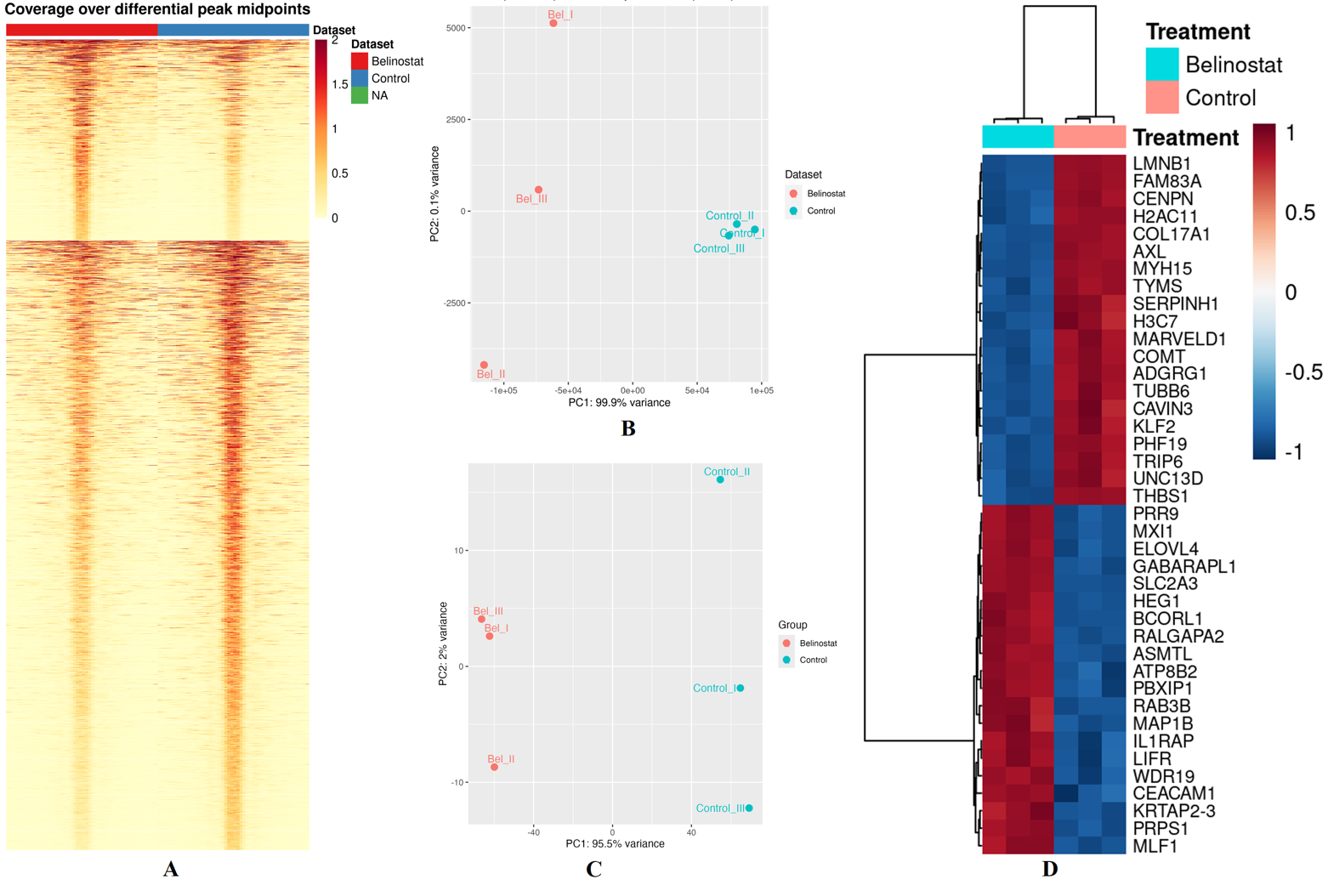

Figure S6: Integrated transcriptomic and epigenomic analysis of belinostat-treated HT-1197 cells. **(A)** Heatmap displaying the mean ATAC-seq signal coverage averaged from three independent replicates centered over identified differential peak midpoints. The center of each sample column corresponds to the midpoint (0) of a 2 Kb window (-1000 to 1000 bp). Rows represent individual differential genomic loci, grouped into distinct clusters based on accessibility patterns. The top cluster shows significantly higher ATAC-seq signal (open chromatin) in the Belinostat-treated group relative to the Control group, while the bottom cluster shows significantly reduced signal (less accessible chromatin) with Belinostat treatment; **(B)** Principal Component Analysis of ATAC-seq data showing control samples and Belinostat-treated replicates; **(C)** Principal Component Analysis of RNA-seq data showing control samples and Belinostat-treated replicates; **(D)** Top 20 up-and down-regulated genes identified by DEseq2 differential gene expression analysis.

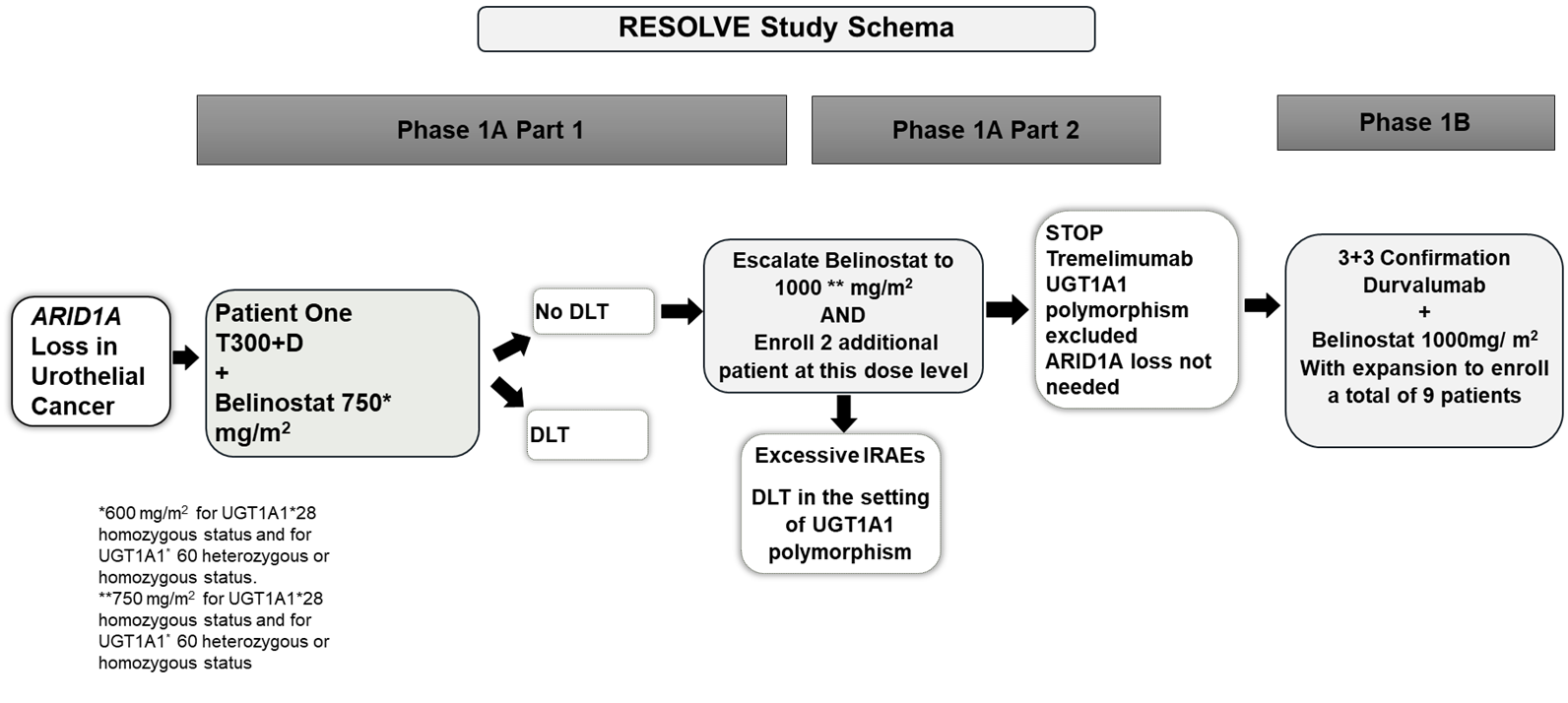

Figure S7: RESOLVE study schema.

Table S1: BAF genes.

| Gene | Chr | BAF type |
| --- | --- | --- |
| *ACTB* | 7p22.1 | Shared |
| *ACTL6A* | 3q26.33 | Shared |
| *ACTL6B* | 7q22.1 | Shared |
| *ARID1A* | 1p36.11 | canonical |
| *ARID1B* | 6q25.3 | canonical |
| *ARID2* | 12q12 | PBAF |
| *BCL11A* | 2p16.1 | canonical |
| *BCL11B* | 14q32.2 | canonical |
| *BCL7A* | 12q24.31 | Shared |
| *BCL7B* | 7q11.23 | Shared |
| *BCL7C* | 16p11.2 | Shared |
| *BICRA* | 19q13.33 | GBAF |
| *BICRAL* | 6p21.1 | GBAF |
| *BRD7* | 16q12.1 | PBAF |
| *BRD9* | 5p15.33 | GBAF |
| *DPF1* | 19q13 | Canonical |
| *DPF2* | 11q13.1 | Canonical |
| *DPF3* | 14q24.2 | Canonical |
| *PBRM1* | 3p21.1 | PBAF |
| *PHF10* | 6q27 | PBAF |
| *SMARCA2* | 9p24.3 | Shared |
| *SMARCA4* | 19p13.2 | Shared |
| *SMARCB1* | 22q11.23 | Shared |
| *SMARCC1* | 3p21.31 | Shared |
| *SMARCC2* | 12q13 | Shared |
| SMARCD1 | 12q13.12 | Shared |
| SMARCD2 | 17q23.3 | Shared |
| SMARCD3 | 7q36.1 | Shared |
| SMARCE1 | 17q21.2 | Shared |
| SS18 | 18q11.2 | Shared |

Note: PBAF, polybromo-associated BAF; GBAF, GLTSCR1-containing BAF, also known as non-canonical BAF; canonical, canonical BAF; shared, subunits shared by at least two BAF variants.

Table S2: Genes demonstrating mutual exclusivity with BAF^m^ identified in ORIEN tumor exomes.

| Gene | p-value |
| --- | --- |
| ***KMT2D*** | 0.00061 |
| *TP53* | 0.0019 |
| ***FGFR3*** | 0.0028 |
| *MAP3K19* | 0.0031 |
| *LYST* | 0.0071 |
| *ZNF302* | 0.0096 |
| *SMAD1* | 0.011 |
| *TBL1XR1* | 0.011 |
| *H2BC3* | 0.011 |
| *EP300* | 0.012 |
| *EHD4* | 0.013 |
| *ESPNL* | 0.013 |
| *DGKZ* | 0.016 |
| *ZNF142* | 0.017 |
| *FHL3* | 0.021 |
| *HECTD3* | 0.021 |
| *LRRC37A3* | 0.021 |
| *ANKZF1* | 0.025 |
| *DLX6* | 0.025 |
| *MX1* | 0.025 |
| *FOXQ1* | 0.026 |
| *ICE1* | 0.027 |
| *SLC39A14* | 0.027 |
| *CLVS2* | 0.027 |
| *PMEL* | 0.027 |
| *TAF4B* | 0.027 |
| *PDE4DIP* | 0.027 |
| *CRYBG3* | 0.033 |
| *RB1* | 0.034 |
| ***KDM6A*** | 0.035 |
| *BTD* | 0.039 |
| *CPSF7* | 0.039 |
| *SOD1* | 0.039 |
| *TCF7* | 0.039 |
| *DPYS* | 0.040 |
| *SPTB* | 0.040 |
| *BTAF1* | 0.041 |
| *GDF10* | 0.042 |
| *GPM6A* | 0.042 |
| *HDAC2* | 0.042 |
| *GNAZ* | 0.044 |
| *MCC* | 0.044 |
| *RRP36* | 0.044 |
| *COL4A2* | 0.044 |
| *SEC24A* | 0.044 |
| *ADARB1* | 0.044 |
| *ALDH6A1* | 0.044 |
| *PDE6B* | 0.044 |
| *RIPK1* | 0.044 |
| *AKT1* | 0.045 |
| *BRIP1* | 0.045 |
| *MAP2* | 0.045 |
| *PCDH17* | 0.045 |
| *PIK3R5* | 0.045 |
| *RASGRP1* | 0.045 |
| *XPC* | 0.045 |
| *COL22A1* | 0.047 |
| *TSC1* | 0.048 |

Note: p-values were not corrected for testing multiple genes. No genes reached statistical significance after multiple-testing correction (Benjamini–Hochberg FDR > 0.05). Genes previously reported to be mutually exclusive with BAF alterations in bladder cancer are highlighted in bold.

Table S3: Pathways demonstrating mutual exclusivity with BAF^m^ identified in ORIEN tumor exomes.

| Pathways | p-value |
| --- | --- |
| GO_REPLICATIVE_SENESCENCE | 0.014 |
| TCGA_GLIOBLASTOMA_MUTATED | 0.018 |
| KEGG_BLADDER_CANCER | 0.020 |
| REACTOME_G_BETA_GAMMA_SIGNALLING_THROUGH_PI3KGAMMA | 0.037 |
| SA_TRKA_RECEPTOR | 0.039 |
| BIOCARTA_G2_PATHWAY | 0.040 |
| BIOCARTA_SPRY_PATHWAY | 0.042 |
| GO_CELL_COMMUNICATION_BY_ELECTRICAL_COUPLING | 0.043 |
| PID_P38_MKK3_6PATHWAY | 0.047 |
| DING_LUNG_CANCER_MUTATED_SIGNIFICANTLY | 0.049 |

Note: p-values were not corrected for testing multiple pathways. No pathways reached statistical significance after multiple-testing correction (Benjamini–Hochberg FDR > 0.05).
